# Evaluating the limitations of Bayesian metabolic control analysis

**DOI:** 10.1101/2025.03.24.644913

**Authors:** Janis Shin, James M. Carothers, Herbert M. Sauro

## Abstract

Bayesian Metabolic Control Analysis (BMCA) has emerged as a promising framework for inferring metabolic control coefficients in data-limited scenarios by integrating Bayesian inference with linlog rate laws. However, its predictive accuracy and limitations remain underexplored. This study systematically evaluates BMCA’s ability to infer elasticity values, flux control coefficients (FCCs), and concentration control coefficients (CCCs) under varying data availability conditions using three synthetic metabolic network models. Our findings highlight the strengths and weaknesses of BMCA, guiding its application in metabolic engineering and emphasizing the need for methodological refinements.

**Author summary:** Understanding how enzymes control metabolic pathways is crucial for optimizing biomanufacturing and synthetic biology applications. Bayesian Metabolic Control Analysis (BMCA) is a promising computational method that integrates Bayesian inference with metabolic control analysis to estimate key control parameters, even in cases with limited experimental data. However, the accuracy and limitations of BMCA remain unclear. In this study, we systematically evaluate BMCA using three synthetic metabolic networks to determine how different types of physiological data impact its predictive performance. We find that BMCA requires flux and enzyme concentration data for accurate predictions, while external metabolite concentrations contribute little. Additionally, BMCA fails to predict elasticity values beyond a magnitude of 1.5 and reliably infer allosteric regulation, even when strong regulatory interactions exist. In addition, BMCA does not accurately rank metabolic control points, which may limit its utility in identifying key enzymes in engineered pathways. Our work provides practical insights into when and how BMCA can be applied, guiding future research in metabolic modeling and control analysis.

## Introduction

Much research has focused on optimizing the process of engineering metabolic pathways to biomanufacture chemicals sustainably [1, 2]. Many groups have demonstrated that it is possible to design alternate routes for carbon molecules through existing metabolic networks in organisms to produce desired chemical products, such as amino acids, oleochemicals, and isoprenoids [3–5]. To meet the demand for biomanufactured chemicals, the process of reengineering endogenous metabolic networks needs to be accelerated.

One approach for reengineering metabolic pathways is to build a kinetic model to identify control points in a metabolic pathway, which can then be targeted to shift metabolic rates in favor of producing the desired product [6–8]. To analyze control distribution within a pathway, some mechanistic models incorporate metabolic control analysis (MCA) [8], a mathematical framework that quantifies how control is distributed among individual enzymes. Using MCA, one can calculate elasticities, which measure how enzyme rates are affected locally by changes in substrate, products, and other effectors. MCA then defines two system-based measurements called flux and concentration control coefficients which are functions of elasticities [9]. The flux control coefficient (FCC) values describe how a change in the concentration of an enzyme can affect fluxes through all the reactions in the pathway. The FCC values with the greatest magnitudes represent the most promising enzyme targets.

While knowledge of the metabolic network structure, allosteric interactions, kinetic parameters, metabolite concentrations, and flux values is valuable for determining control coefficients using MCA, many kinetic parameters within a metabolic pathway remain unknown, necessitating empirical approaches for estimating these coefficients. One method involves measuring the resulting fluxes and concentrations from direct perturbations to the enzyme or perturbing enzymes using inhibitors [10].

Unfortunately, this method does not easily scale. If the metabolite concentrations are known, control coefficients can also be estimated based on elasticities using various techniques, such as conducting simulations to build models (primarily for smaller pathway segments) and performing modulation experiments to assess changes in metabolite levels after manipulating upstream or downstream components. While these methods can provide valuable insights, they do not easily scale to whole-cell studies.

Bayesian Metabolic Control Analysis (BMCA) is an approach to a mechanistic model that combines Bayesian inference with linlog rate laws and whole-cell physiological data to infer elasticity values, which can subsequently be used to estimate control coefficients [11, 12]. This approach is particularly advantageous in data-limited situations, as Bayesian inference allows the incorporation of prior information about a system into the analysis. By leveraging existing information, BMCA enhances the statistical power to detect meaningful associations and effects. In practice, to apply Bayesian inference, one needs a prior, a likelihood function, and observed data, if available. BMCA models each reaction using linlog kinetics [11–13], which are simplified and approximate rate laws made using a steady-state assumption that use elasticity values as parameters—the linlog rate laws function as the likelihood in the Bayesian inference. Although linlog rate laws are an approximation, they appear to be robust to large perturbations to the system [14]. If available, whole-cell data (such as flux values, enzyme concentrations, and metabolite concentrations) can be input into the Bayesian framework as observed data. Through the combination of mechanistic understanding and prior information, BMCA generates posterior probability distributions for the elasticities, from which control coefficients are subsequently estimated.

This paper considers the following questions: What type of physiological data and how much data is necessary for BMCA to make helpful predictions? How effective is BMCA at predicting allosteric interactions? We address these questions using three synthetic models from which we establish ground truth values such as elasticities and control coefficient. We evaluated how accurately BMCA inferred elasticity values across all the reactions in the tested network topologies and FCC rank orders for the reaction directly involved in producing a specific output.

We found that flux data, enzyme concentration data, and internal metabolite concentration data were critical for determining elasticity values and concentration control coefficients, whereas flux data and enzyme concentration data were more critical for determining flux control coefficients. Contrary to previous claims about the BMCA algorithm’s ability to detect implicit allosteric relationships [11], our work demonstrated that BMCA failed to detect allosteric relationships for the three models we investigated, even if the allostery was strong. Additionally, the BMCA prior predictions and posterior predictions for the top ten reactions with the highest FCCs were only marginally different. These insights can provide future guidance for researchers when using BMCA and help clarify the predictive limits of BMCA.

## Building the test models and simulated datasets

Three different model topologies were constructed (Fig. 1), two of which included variations incorporating allosteric regulators, for a total of seven model variations. Topology A (TopA) is a linear chain, with several branch points. TopA has three total variations of possible allosteric regulation: one without any, one with metabolite J regulating the reaction labeled v6, and one with metabolite J regulating reaction v6 in addition to metabolite G regulating reaction v3. For all TopA models, reaction v15 was optimized. Topology B (TopB) exhibits more branching than TopA and also has three total variations of possible allosteric regulation: one without any, one with metabolite H regulating reaction v5, and one with metabolite H regulating reaction v5 in addition to metabolite O regulating reaction v14. Topology C is loosely based on E. coli central metabolism and contains not only conserved moieties (which Topology A and B do not have), but also a cycle in the form of the Krebs cycle. Summaries of ground truth coefficient values for all three networks can be found in (Table 1).

**Table 1.** Table Summaries of ground truth coefficient value ranges for all three networks and their allosteric variations. The first number is the minimum value and the second is the maximum value.

|  |  | noReg | Reg1 | Reg2 | Many regulators |
| --- | --- | --- | --- | --- | --- |
| TopA | elasticities | -1.43 / 1.45 | -1.58 / 1.85 | -1.70 / 1.71 | - |
|  | CCCs | 0.84 / -1.36 | 0.78 / -1.14 | 0.74 / -0.98 | - |
|  | FCCs | 0.96 / -0.83 | 0.96 / -0.78 | 0.95 / -0.79 | - |
| TopB | elasticities | -10.94 / 11.74 | -14.95 / 15.75 | -22.82 / 23.64 | - |
|  | CCCs | 2.53 / -1.7 | 1.87 / -1.6 | 1.86 / -1.60 | - |
|  | FCCs | 1.15 / -0.86 | 1.03 / -0.96 | 1.02 / -0.96 | - |
| TopC | elasticities | - | - | - | -1.8E5 / 1.8E5 |
|  | CCCs | - | - | - | 2.07 / -2.01 |
|  | FCCs | - | - | - | 5.93 / 3.21 |

**Fig 1.**
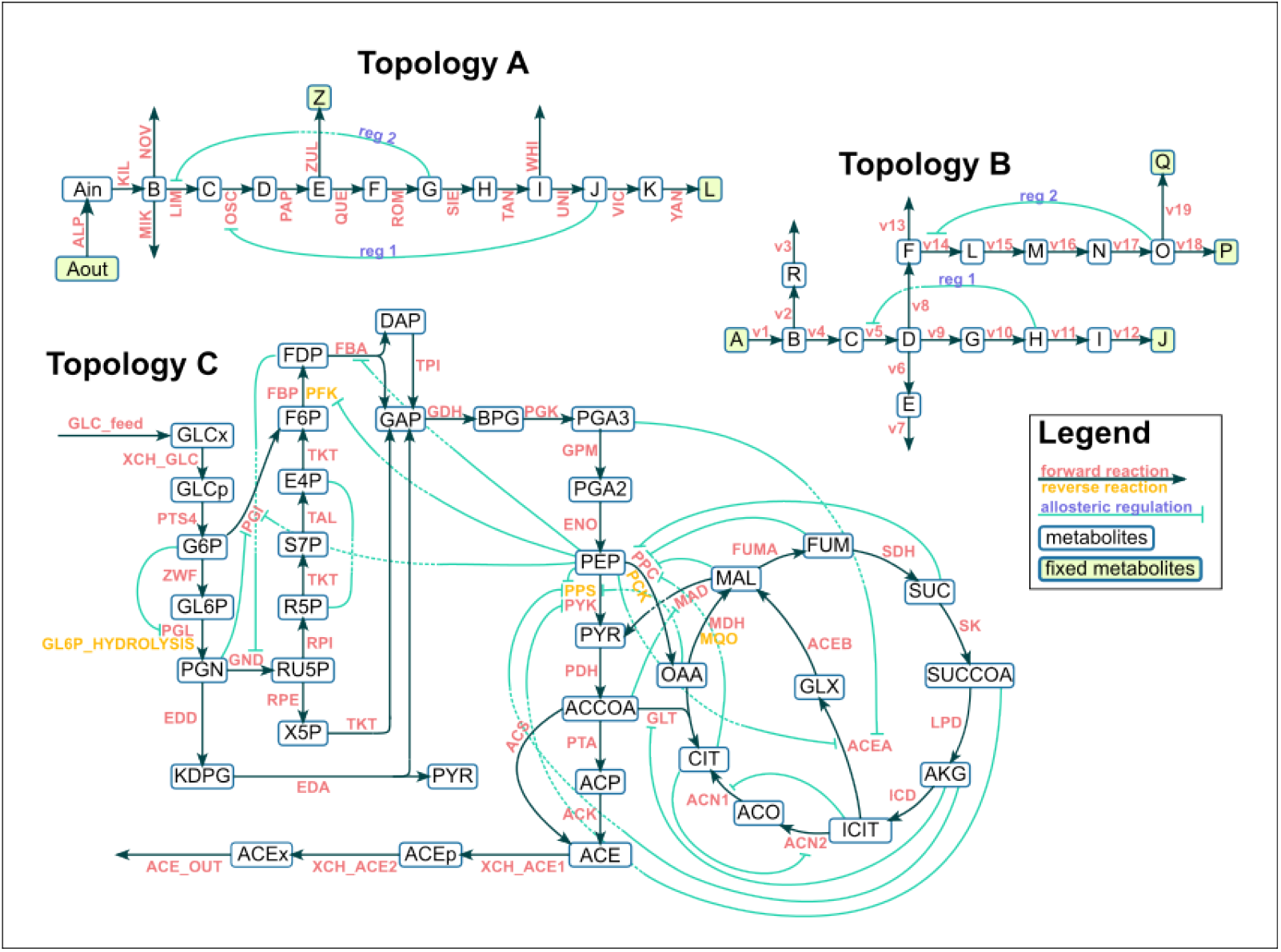
Three synthetic networks used to benchmark BMCA. A) Topology A, B) Topology B, C) Topology C, adapted from Millard et al (15). Cyan lines describe the allosteric regulation. For Topology C, only negative allosteric regulation is shown.

## Materials and methods

### Construction of test models

TopA was based on a yeast glycolysis model by Teusink et al. (BIOMD0000000064) [10, 15]. The ATP and NADH assignment rules were removed and the enzyme parameters were changed to values so that the reactions were not at equilibrium. TopA has 14 metabolites and 16 reactions Reaction v15 is the reaction being optimized in all the BMCA analyses. TopB has 17 metabolites and 19 reactions. Reaction v19 is the reaction being optimized in all the BMCA analyses. TopC was based on an E. coli model by Millard et al. (MODEL1505110000) [16] that includes the pentose phosphate pathway and the TCA cycle. It has 64 metabolites and 68 reactions with the flux through reaction ACE OUT to be optimized.

### Generating simulated data

Models and their variations were run to steady state using Tellurium [17, 18] for the following enzyme perturbation levels: 10%, 20%, 30%, 40%, 50%, 150%, 300%, 500%, 700%, and 1000%. At steady state, the relative enzyme concentrations and the absolute metabolite concentration and reaction fluxes were recorded. Then, each enzyme was perturbed by a set perturbation level (e.g. 10% of the original enzyme level) and the absolute steady state metabolite concentrations and reaction fluxes were recorded as was the relative enzyme concentrations for all enzymes. The model was reset after each enzyme perturbation so that the generated data only reflected the results of one enzyme being perturbed. If steady state could not be reached at a given enzyme concentration, the data for that perturbation were excluded from the final datasets.

### Running BMCA

When all data are provided, the linlog model is formulated as:

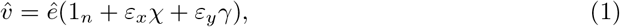

where 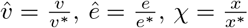, and 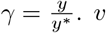 represents the flux data, *e* the enzyme concentrations, *x* the internal metabolite concentrations, and *y* the external metabolite concentrations. The star denotes values at the reference state. Any experiment can be chosen as the reference state. For this paper, the reference state was designated as the experiment where none of the enzymes were perturbed, for simplicity.

Elasticity parameters *ε*_*x*_ and *ε*_*y*_ are initialized as deterministic transformations. The normalized data corresponding to the observed variables (*ê*, *χ*, and *γ*) are modeled as normal distributions. The likelihood function (Eq. 1) represents the expected flux values based on these observations. Bayesian inference is performed using Automatic Differentiation Variational Inference (ADVI), which optimizes the evidence lower bound (ELBO) to approximate the posterior distributions of the model parameters.

When specific data types are omitted, the missing values are treated as unmeasured data, and latent variables are introduced to account for the missing information. In cases where flux or enzyme data are unavailable, the posterior distributions of these latent variables are later used to estimate Control Coefficients (CCCs) and Flux Control Coefficients (FCCs). If flux data are omitted, only the reference fluxes are supplied to the Bayesian inference model, and the linlog model is reformulated to solve for enzyme concentrations instead of fluxes as follows:

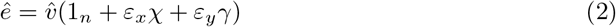

### Analyses

The CCCs and FCCs are estimated using the method described by St. John et al. [11]. However, FCC predictions for fluxes in which an enzyme perturbs its own reaction exhibited a consistent deviation of -2. To account for this systematic offset, a correction factor of +2 was applied to these predictions S1 Fig.

## Results

### Data type omission experiments

The goal of this paper is to determine which data types are most critical for BMCA predictions and whether BMCA can accurately rank flux control coefficients (FCC), even if the values themselves are uncertain. To address these questions, we tested BMCA using all model topologies and datasets, with outcomes analyzed across different perturbation and allostery levels and proportions of observed data input. In our study, we evaluated each model variation using the following experimental setup (Fig. 2).

**Fig 2.**
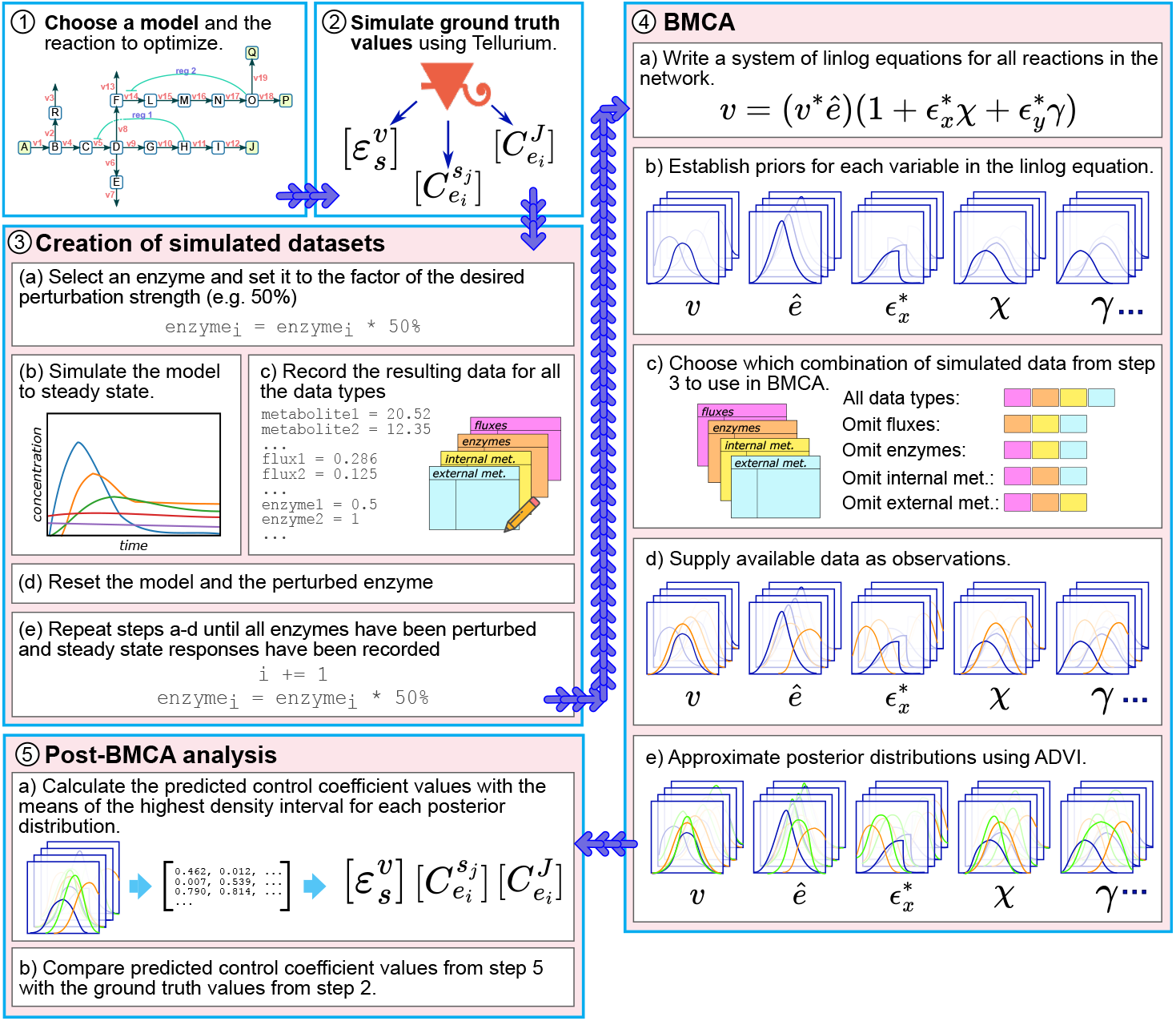
Overview of BMCA benchmarking experiments. Synthetic models are simulated using tellurium (16) to produce datasets of fluxes and concentrations of enzymes, internal metabolites, and external metabolites. The ground truth FCC values and elasticities are also calculated from the simulation. For each model variation, five parallel BMCA runs take place: one without any omitted data and one each of omitting one data type. Each BMCA run results in a set of posterior distributions for elasticities. The HDI for each posterior distribution is computed, resulting in a single value for each elasticity. These elasticity sets are compared with ground truth elasticity values and then used to calculate control coefficient values which are subsequently compared with the ground truth control coefficients.

Ground truth values for the elasticities and control coefficients were calculated for each model. The models were then simulated to steady state and four types of datasets were generated: flux values for each reaction and concentrations for each enzyme, internal metabolite, and external metabolite. The choice of data types to include or omit from the BMCA algorithm as well as the presence of the allosteric regulators was varied.

Each variation was used as the observational data for the BMCA algorithm. The variation where no data was omitted represented a best-case scenario for the study. We then executed BMCA multiple times, omitting one data type at a time to assess the contribution of each type to the predictive capability of the algorithm. Each run produced posterior distributions for all elasticities, from which we calculated the mean of the highest density interval (HDI). These means were compared against the ground truth values, and FCCs were estimated from the elasticity HDI means. Finally, we ranked the reactions based on both ground truth and predicted FCC values for comparison.

### Omitting different data types leads to different BMCA elasticity predictions

Elasticity values generated by BMCA in various conditions and model variations were compared to ground truth to identify factors that influenced the BMCA algorithm’s performance. The mean for each HDI from each posterior distribution for each elasticity was calculated for comparison. In Topology A, when BMCA was given all the data, the predicted elasticity values matched the ground truth values well, even for variations with allosteric regulation (Fig. 3). The absence of internal metabolite concentrations led BMCA to predict nearly all elasticities as zero. When enzyme concentrations were omitted, elasticities with negative ground truth values were predicted close to zero, while those with positive ground truth values were underestimated. Conversely, omitting external metabolite data had minimal impact on the algorithm’s performance, as the results remained nearly identical to those obtained with all data types included. Additionally, despite not receiving flux data, the BMCA algorithm was still able to determine whether an elasticity is negative or positive; however, it resulted in the majority of predicted elasticities being either 1 or -1. The same patterns were also observed in Topology B (Fig. 4). However, in Topology B, only the elasticities whose ground truth values were within -2 and 2 were predicted well when the BMCA algorithm was given all the data or when the external metabolite concentrations were withheld from the algorithm. The elasticities outside of that range were not predicted well.

**Fig 3.**
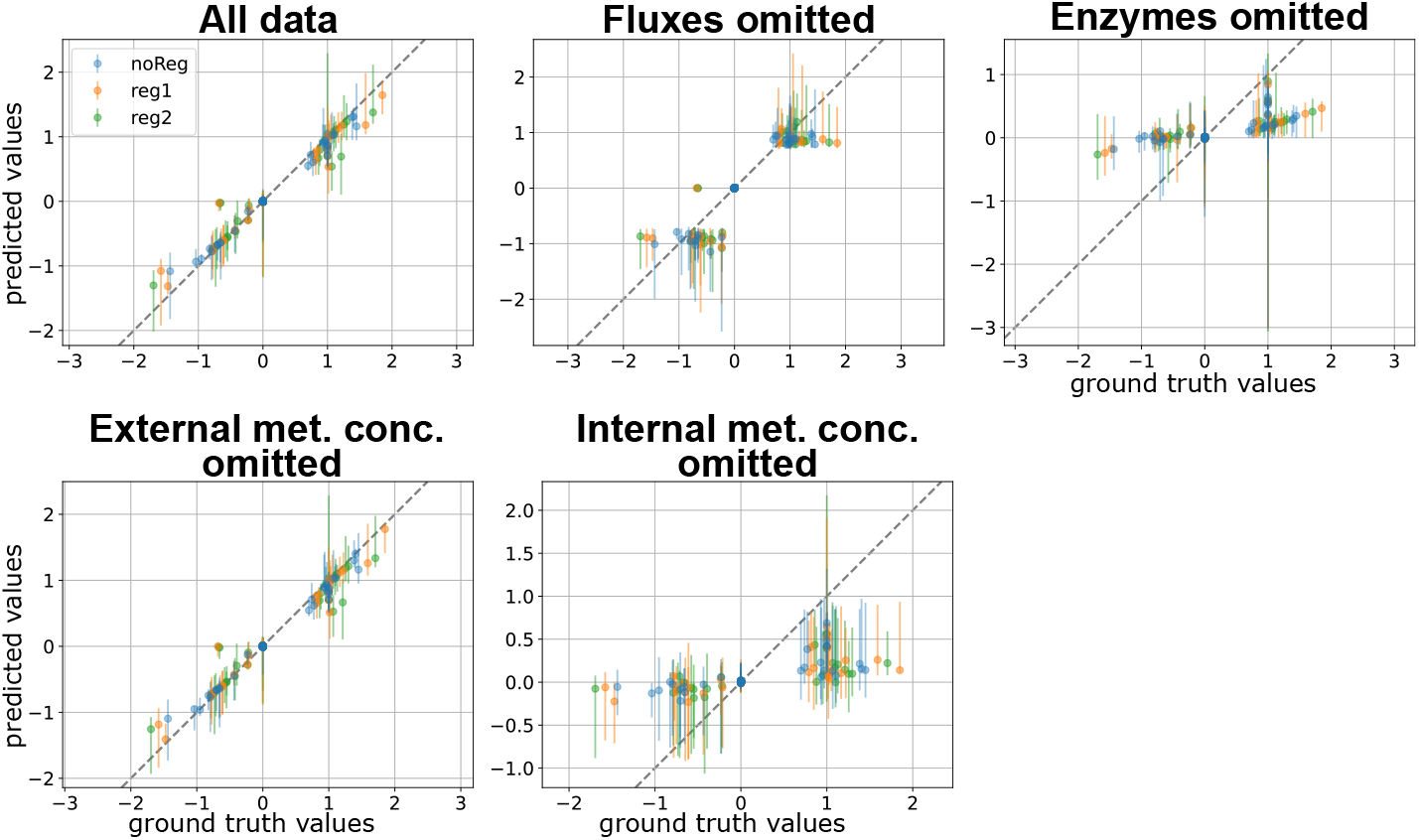
BMCA elasticity predictions compared against ground truth values for Topology A. Each dot signifies the median while the error bars represent the range of the predicted elasticity values across the different enzyme perturbation levels tested. The titles for each graph indicate which data type was omitted when running BMCA.

**Fig 4.**
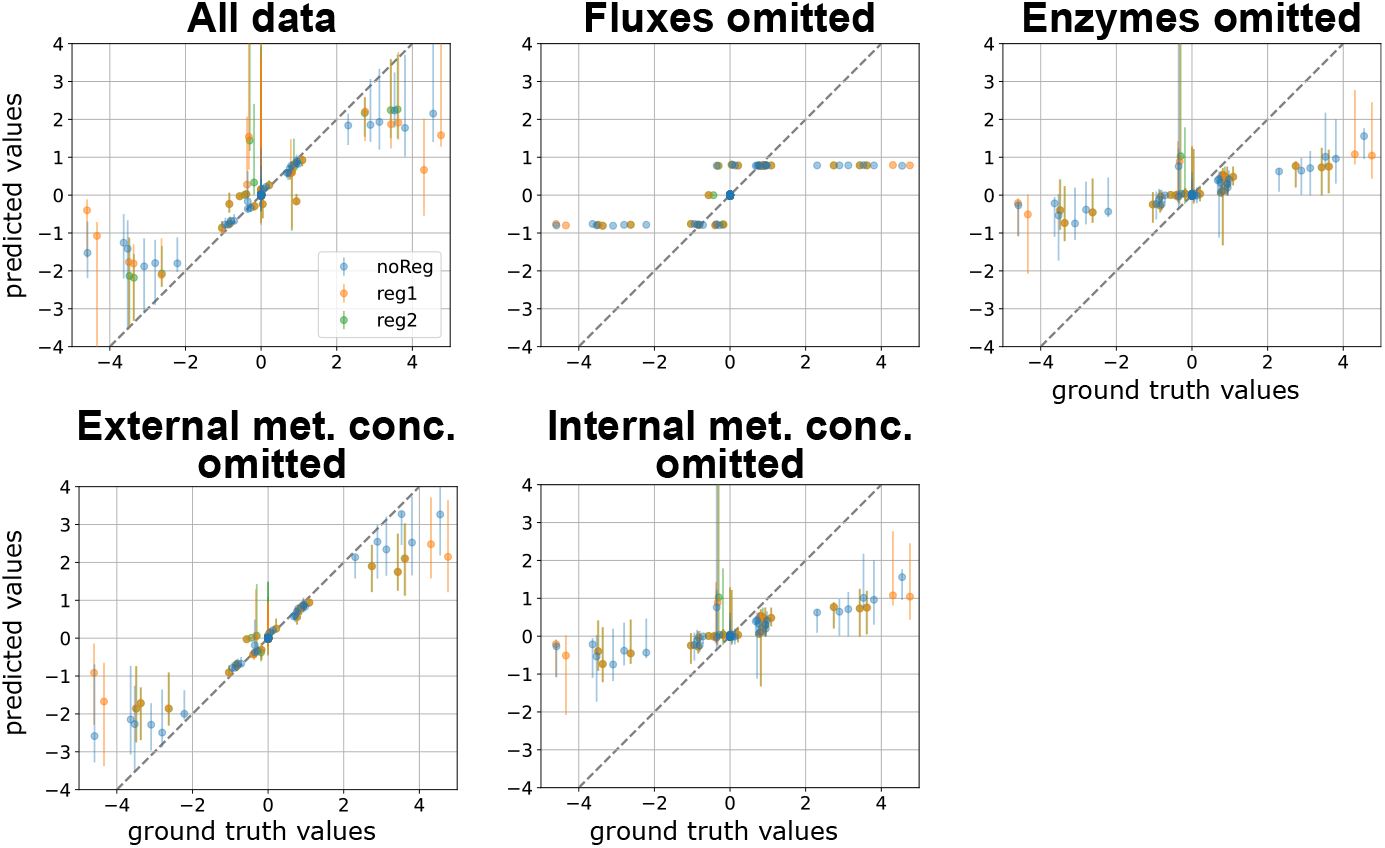
BMCA elasticity predictions compared against ground truth values for Topology B. Each dot signifies the median while the error bars represent the range of the predicted elasticity values across the different enzyme perturbation levels tested. The titles for each graph indicate which data type was omitted when running BMCA. All of the graphs are zoomed in as the ground truth elasticity.

### Ground truth elasticities with magnitudes larger than 1.5 are often underestimated

BMCA seems to demonstrate a range for which it can make good predictions for elasticity values. For all cases of providing the BMCA algorithm with all the data, the expectation is that the algorithm should predict the elasticity values correctly. This is observed in the elasticity prediction results for Topology A when no data was omitted. However, providing complete data to the BMCA algorithm did not produce perfect elasticity predictions for Topology B and C. The BMCA algorithm predicts well for a small range of values between -1.5 and 1.5. Since the ground truth elasticity values for Topology A are small, the BMCA algorithm has no problem predicting all of its elasticity values. In contrast, Topologies B and C have much larger ground truth elasticity values and predictions for the ground truth elasticity values outside of the -1.5 and 1.5 range are capped at about -2 and 2 (Fig. 4, 5).

**Fig 5.**
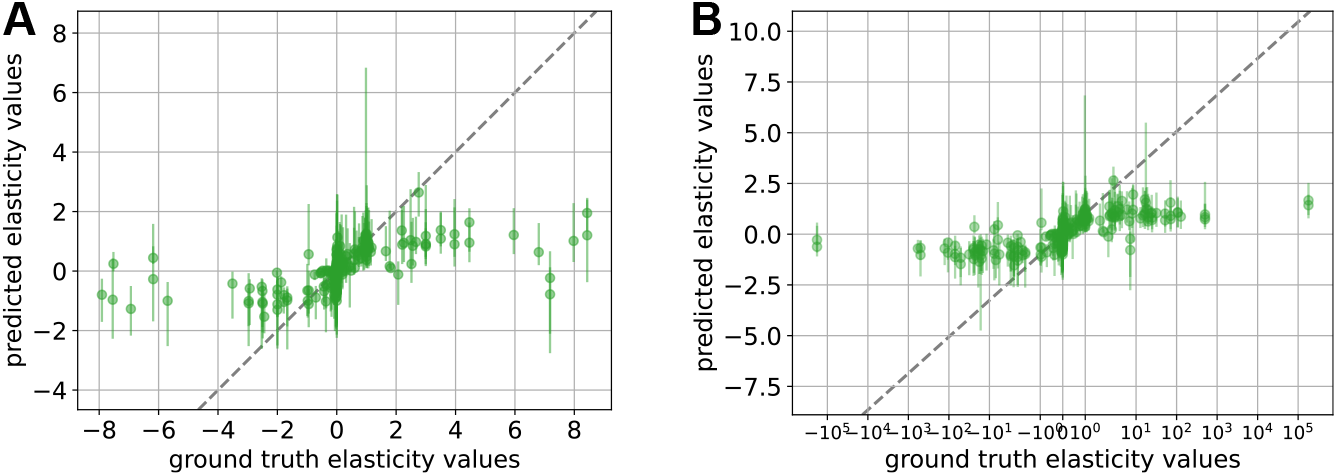
BMCA-predicted elasticities for Topology C across all perturbation strengths compared against ground truth values. A) Comparison of elasticities for ground truth values with less than a magnitude of 10. B) Comparison of elasticities for all ground truth values. Each dot signifies the median while the error bars represent the range of the predicted elasticity values across the different enzyme perturbation levels tested.

### Effect of varying perturbation levels and regulator presence on elasticity predictions across network variations

Levene’s test was used to determine whether elasticity predictions differed in variance across perturbation levels and varying degrees of allostery. This test assesses whether the groups being compared have equal variances. Variability in the data was analyzed instead of central tendency (ANOVA or Kruskal-Wallis test), as all sets of elasticity predictions, regardless of perturbation or allostery level, have a mean and median of zero. Analysis of the BMCA results revealed that the algorithm’s elasticity predictions remained largely unaffected by variations in enzyme perturbation levels for TopA (Table 2). In contrast, for TopB variations with allosteric regulators, perturbation strength significantly influenced the variance of elasticity predictions, as indicated by Levene’s test. For TopC, which contains numerous regulators, elasticity predictions varied significantly across all perturbation levels.

**Table 2.** Results of Levene’s test comparing predicted elasticities across varying perturbation levels for different network variations. The first value represents the Levene test statistic, while the second value denotes the corresponding p-value. Comparisons where no significant difference in perturbation strengths was detected are highlighted in blue, while those indicating significant differences are highlighted in red. All values are rounded to three decimal places.

|  | noReg | Reg1 | Reg2 | Reg1-stronger | Reg2-stronger | Many regulators |
| --- | --- | --- | --- | --- | --- | --- |
| TopA | 8.30e-2 / 9.99e-1 | 1.74e-1/ 9.97e-1 | 2.90e-1 / 9.78e-1 | 4.35e-1 / 9.17e-1 | 3.41e-1 / 9.61e-1 | - |
| TopB | 1.04e-1 / 9.99e-1 | 4.60 / 4.76e-6 | 4.23 / 1.86e-5 | 4.25 / 1.76e-5 | 5.31 / 3.30e-7 | - |
| TopC | - | - | - | - | - | 1.52e+01 / 4.93e-25 |

However, when comparing elasticity predictions across different regulatory conditions (e.g. TopB-noReg, TopB-reg1, and TopB-reg2) at the same perturbation levels, no significant differences were observed (Table 3).

**Table 3.** Results of Levene’s test comparing the predicted elasticities under varying levels of regulation for each perturbation level in networks TopA and TopB with no regulation. The top value represents the Levene test statistic, while the bottom value denotes the corresponding p-value. Comparisons where no significant difference in perturbation strengths was detected are highlighted in blue.

|  | 0.1x | 0.2x | 0.3x | 0.4x | 0.5x | 1.5x | 3x | 5x | 7x | 10x |
| --- | --- | --- | --- | --- | --- | --- | --- | --- | --- | --- |
| TopA | 0.01 | 0.02 | 0.01 | 0.02 | 0.01 | 0.02 | 0.05 | 0.06 | 0.06 | 0.06 |
|  | 0.99 | 0.98 | 0.99 | 0.98 | 0.99 | 0.98 | 0.95 | 0.94 | 0.94 | 0.94 |
| TopB | 0.46 | 0.17 | 0.17 | 0.16 | 0.12 | 0.28 | 0.08 | 0.09 | 0.20 | 0.39 |
|  | 0.63 | 0.85 | 0.84 | 0.85 | 0.88 | 0.75 | 0.92 | 0.91 | 0.82 | 0.67 |

### Comparison of BMCA CCC predictions with ground truth values

The BMCA algorithm’s predictions of CCCs for Topology A aligned with its elasticity predictions in several ways (Fig. 6). When provided with all data, predictions were accurate. Adding allosteric regulation reduced precision, while omitting fluxes generally preserved the correct sign of ground truth values but produced inconsistent and uninformative results otherwise. Omitting enzymes and internal metabolites caused overestimation, particularly for ground truth CCCs near zero. Excluding external metabolites had minimal impact, yielding results similar to using the full dataset.

**Fig 6.**
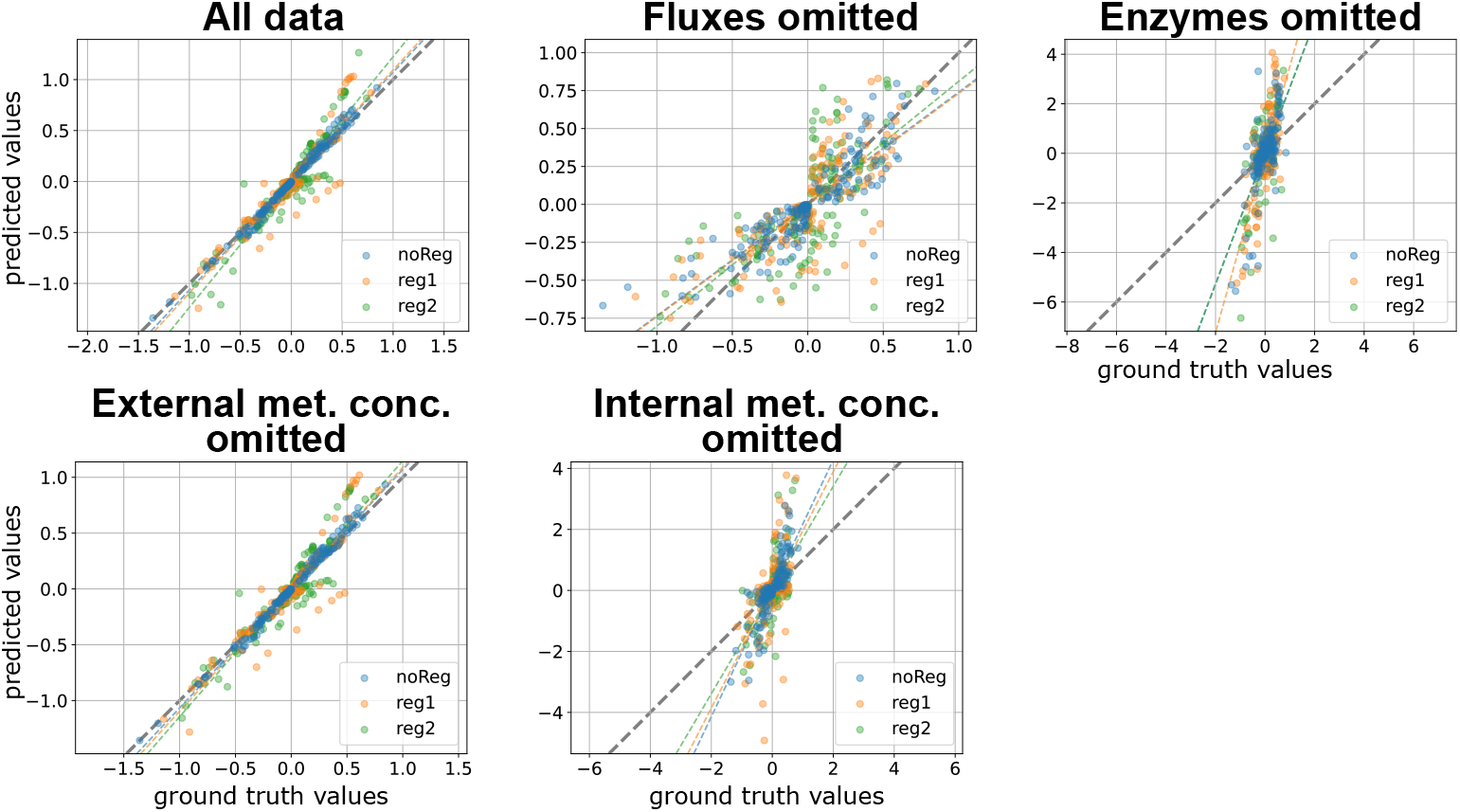
BMCA CCC predictions compared against ground truth values for Topology A. A) Topology A, B) Topology B, C) Topology C, adapted from Millard et al [16]. Cyan lines describe the allosteric regulation.

For Topology B, similar trends emerged but on a larger scale (Fig. 7). When no data was omitted, predictions matched ground truth values closely, particularly in datasets without regulators. However, datasets implying regulator presence introduced greater variance in predictions (Fig. 8). Notably, excluding external metabolites did not degrade the BMCA algorithm’s predictive accuracy, even with the datasets that demonstrated implicit allosteric regulation. As with Topology A, omitting fluxes preserved the sign of ground truth values but resulted in poor predictions while omitting enzymes and internal metabolites led to significant overestimation of CCCs.

**Fig 7.**
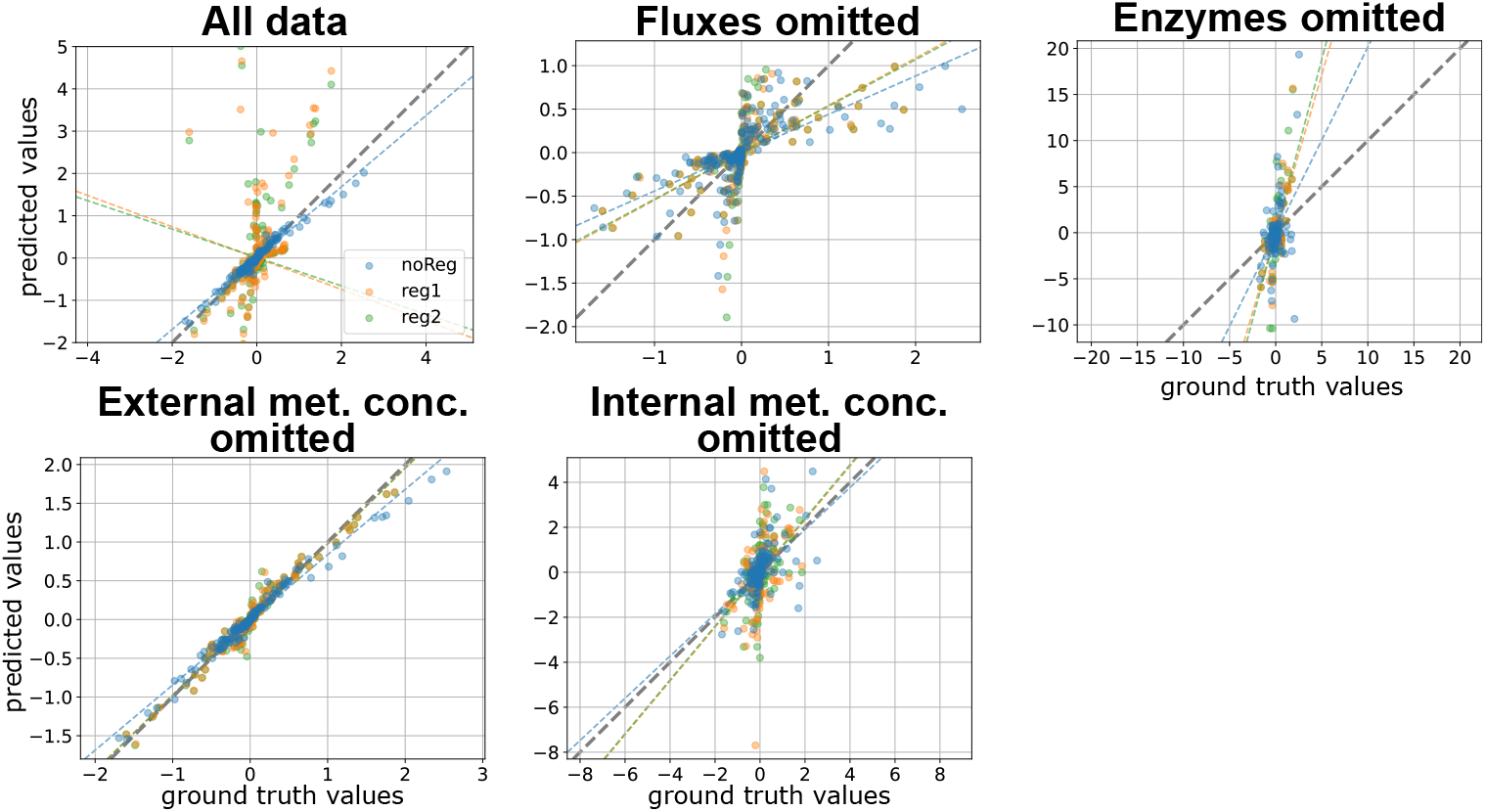
BMCA CCC predictions compared against ground truth values for Topology B. Each dot signifies the median of the predicted CCC values for the different enzyme perturbation levels tested. The titles for each graph indicate which data type was omitted when running BMCA.

**Fig 8.**
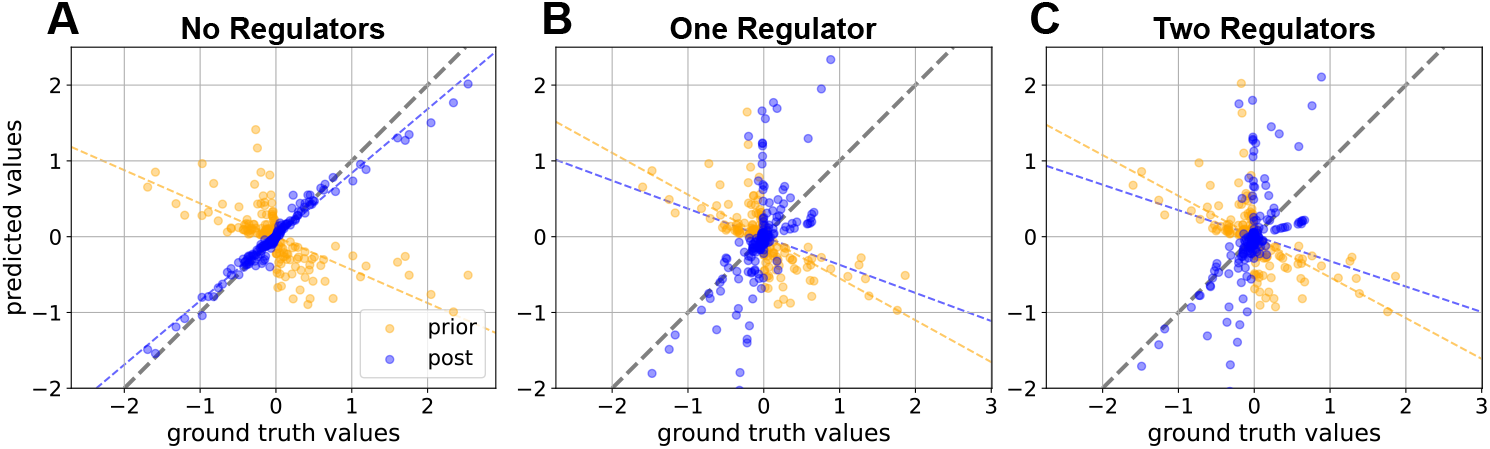
BMCA CCC predictions compared against ground truth values for Topology B model variations when no data types were omitted. A) no regulators, B) one regulator, and C) two regulators. These graphs are zoomed into the main mass of dots and do not include each dot. Each dot signifies the median of the predicted CCC values for the different enzyme perturbation levels tested.

Although the BMCA algorithm was provided with the complete dataset for Topology C, it produced numerous nonzero CCC values for elasticity where the ground truth values were zero, and vice versa (Fig. 9). The ground truth CCC values ranged from just above 2 to just below -2, with the predictions remaining within these bounds. However, the uncertainty ranges were large—sometimes exceeding ten times the predicted value—rendering them uninformative.

**Fig 9.**
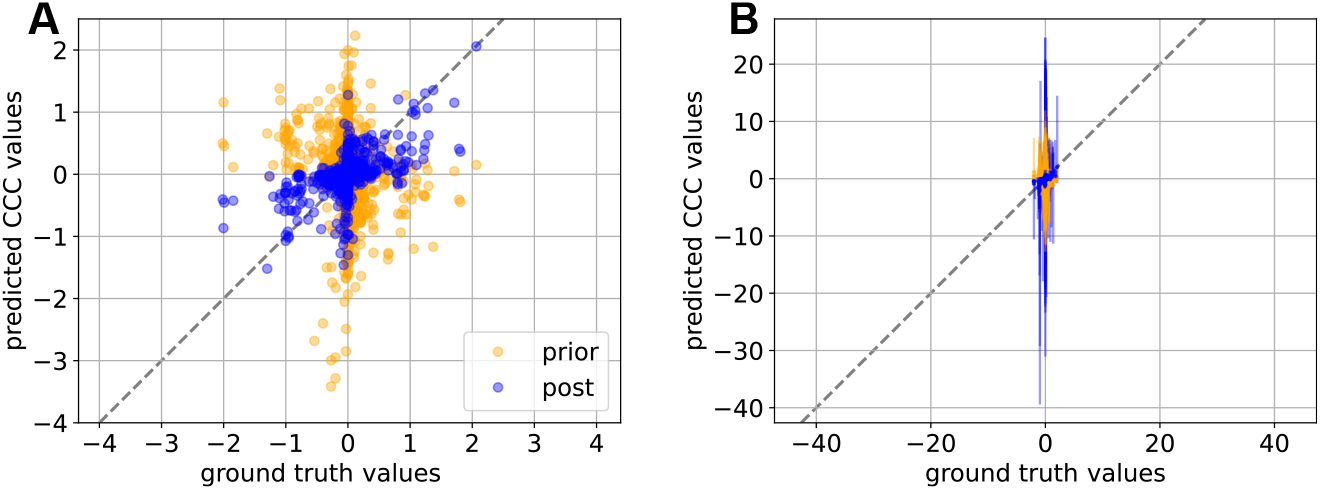
CCC value comparisons for Topology C. A) Median of CCC values calculated from all perturbations of HDI elasticity distributions predicted by BMCA. B) Ranges of possible values from the different perturbations.

### Comparison of FCC predictions with ground truth values

The omission of various data types significantly impacted the FCC predictions made by the BMCA algorithm (Fig. 10). In Topology A, no discernible difference was observed between providing all data and omitting external metabolite concentration data, suggesting that the addition of external metabolite concentration data does not make a difference to BMCA FCC predictions at this scale. Similarly, omitting internal metabolite concentrations resulted in slightly more variance in the FCC predictions, although the overall effect was minimal. The exclusion of enzyme data resulted in some instances of FCC overestimation, although the majority of the data points were predicted within a 0.1 margin of their respective ground truth values.

**Fig 10.**
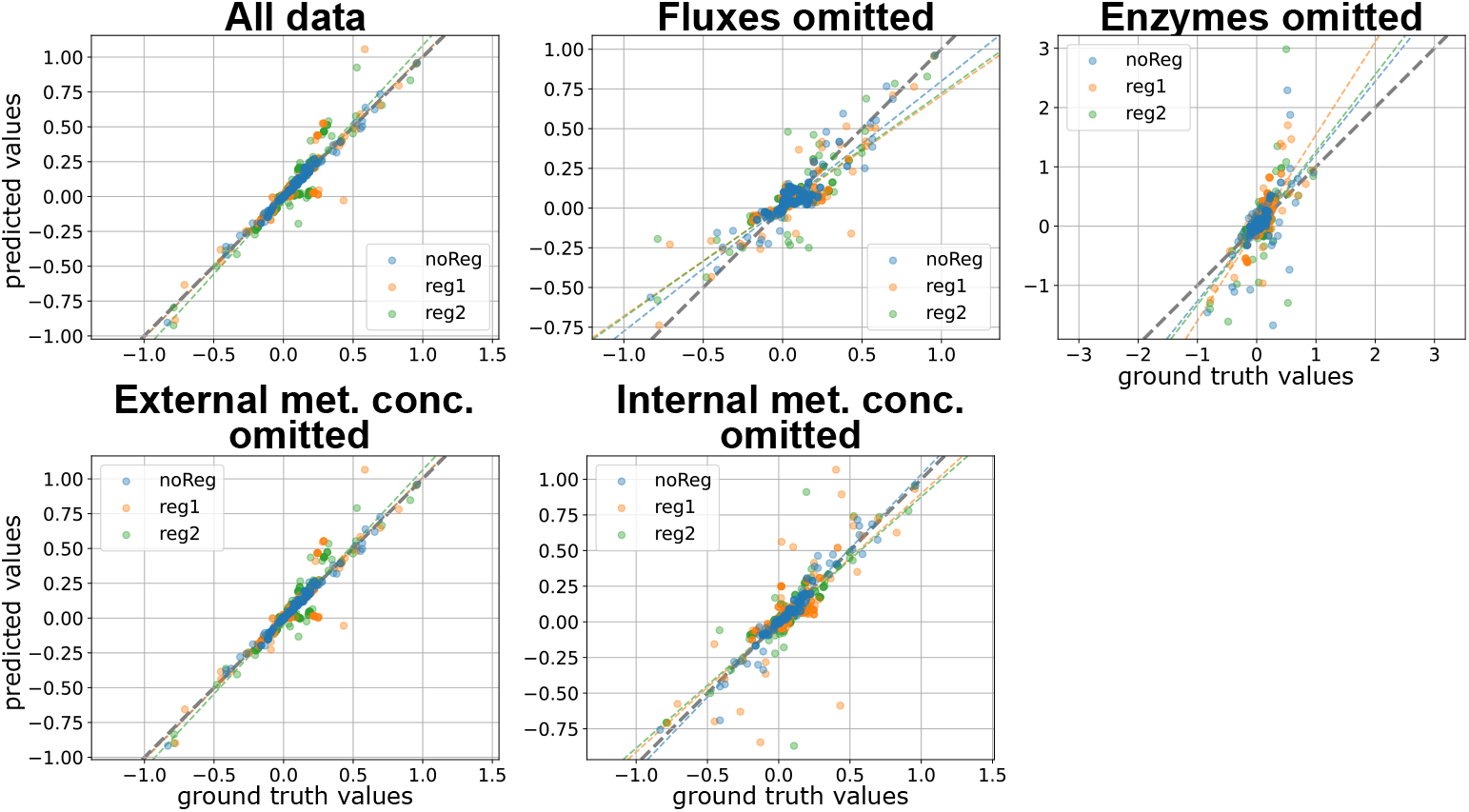
BMCA FCC predictions compared against ground truth values for Topology A. Each dot signifies the median of the predicted elasticity values across the different enzyme perturbation levels tested. The titles for each graph indicate which data type was omitted when running BMCA. Corrections for FCC values for reactions whose enzymes are being perturbed have been applied.

The uncertainty ranges for the FCC predictions were uninformative and varied wildly when there were no data omissions or when enzyme data or internal metabolite concentration data was withheld from the BMCA algorithm (Fig. 11). When flux data was omitted, the resulting uncertainty ranges did not cover the actual ground truth FCC values. Omitting only external metabolite concentrations led to reasonable uncertainty ranges that included the ground truth FCC values.

**Fig 11.**
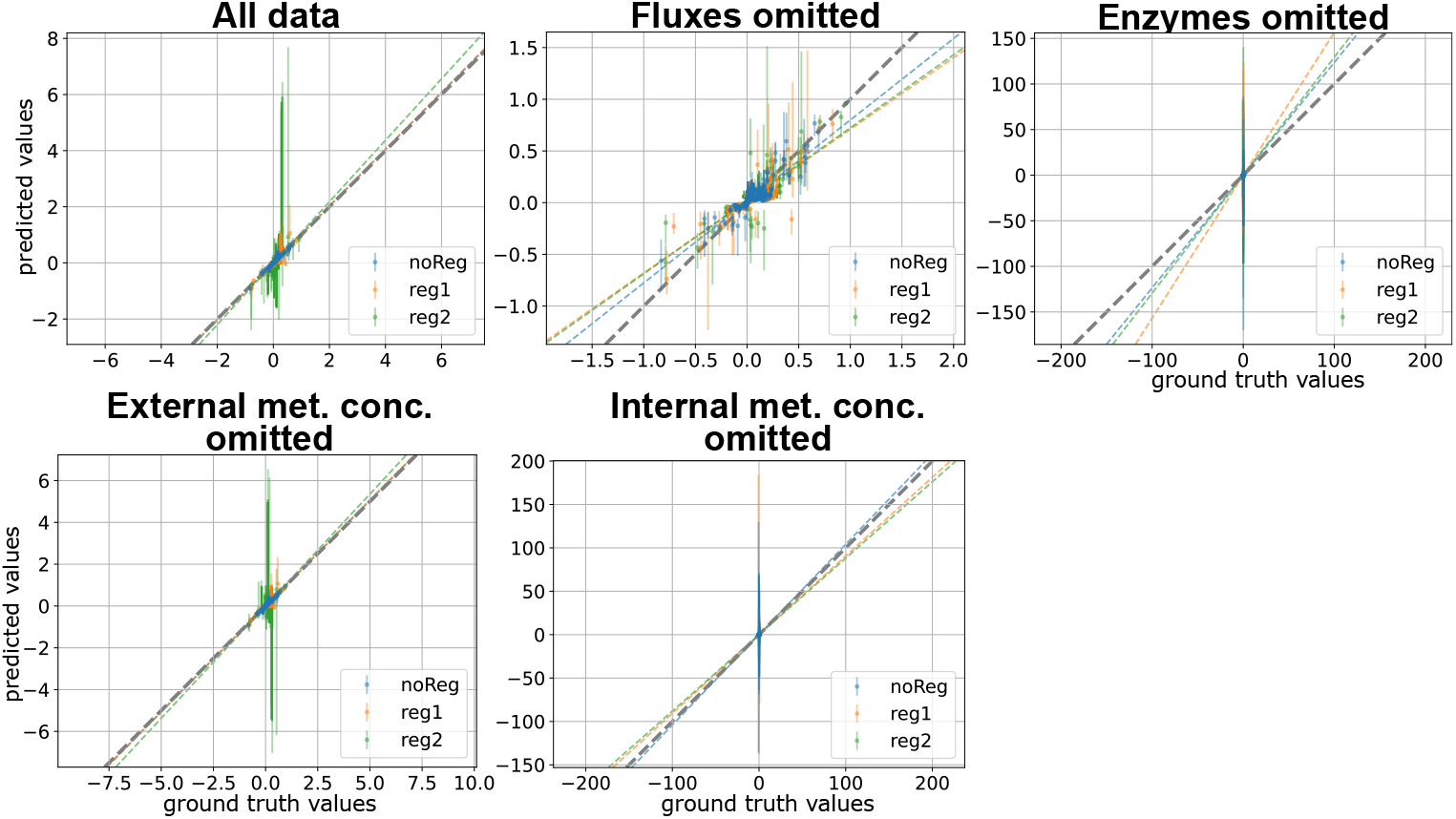
Uncertainty ranges for BMCA FCC predictions compared against ground truth values for Topology A. Each dot signifies the median while the error bars represent the range of the predicted elasticity values across the different enzyme perturbation levels tested. The titles for each graph indicate which data type was omitted when running BMCA. Corrections for FCC values for reactions whose enzymes are being perturbed have been applied.

Excluding flux data had a more pronounced impact, leading to decreased precision in the predicted FCC values, particularly as the ground truth values deviated further from zero. BMCA both underestimated and overestimated the magnitude of FCC values under these conditions, and the omission of flux data yielded little to no change between the prior and posterior distributions (Fig. 12). These results indicate that flux data is essential for accurate FCC predictions.

**Fig 12.**
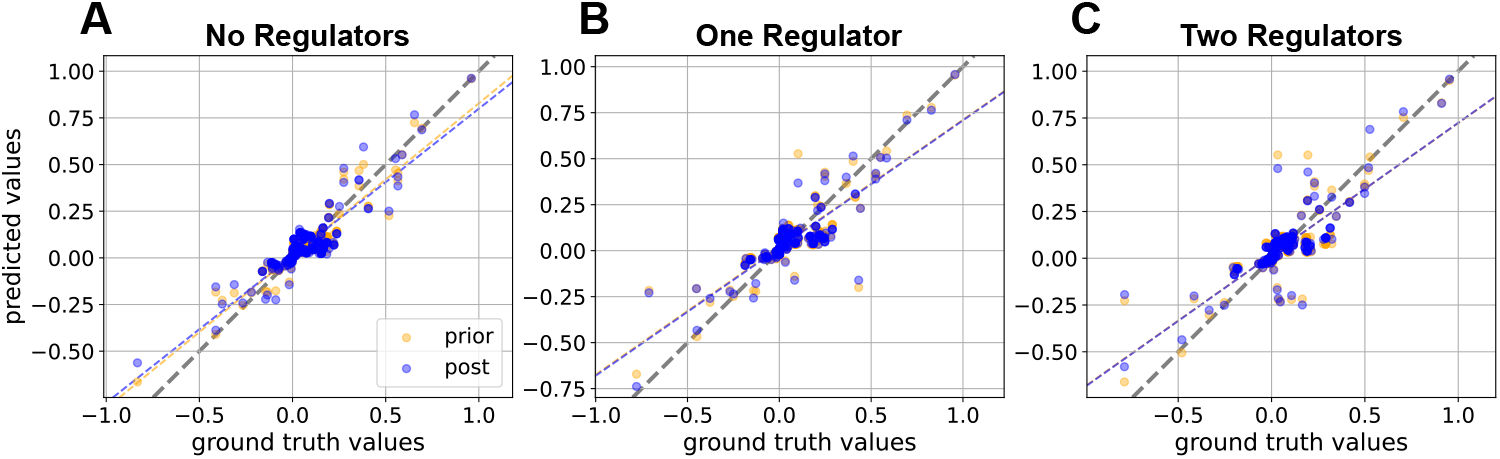
BMCA FCC predictions compared against ground truth values for Topology A model variations when flux values were omitted. A) no regulators, B) one regulator, C) two regulators. Each dot signifies the median while the error bars represent the range of the predicted elasticity values across the different enzyme perturbation levels tested. The titles for each graph indicate which data type was omitted when running BMCA. Corrections for FCC values for reactions whose enzymes are being perturbed have been applied.

For Topology B, which is characterized by a more branched network, the FCC prediction behaviors observed in the omission experiments in Topology A were similar but at a larger scale (Fig. 13). Supplying the BMCA algorithm with all available data in the absence of allosteric regulation resulted in posterior predictions close to their respective ground truth values. However, model variations with allosteric regulation exhibited some overestimation in FCC values. Interestingly, excluding external metabolite data seemed to perform better than using all the data, especially for model variations that included allosteric regulators.

**Fig 13.**
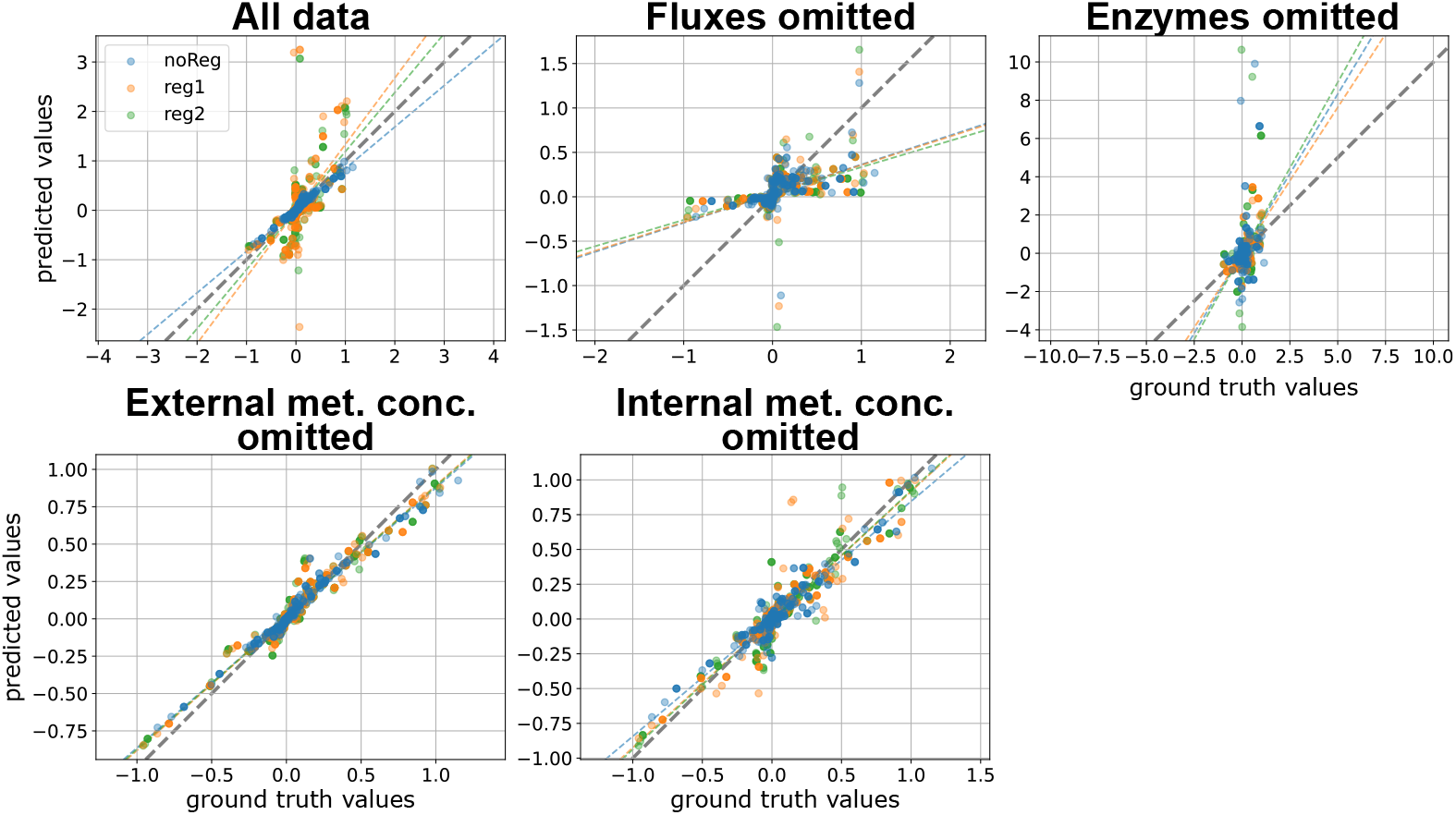
BMCA FCC predictions compared against ground truth values for Topology B. Each dot signifies the median while the error bars represent the range of the predicted elasticity values across the different enzyme perturbation levels tested. The titles for each graph indicate which data type was omitted when running BMCA. Corrections for FCC values for reactions whose enzymes are being perturbed have been applied.

Excluding flux data in Topology B resulted in little difference between the prior and posterior FCC predictions (Fig. 14), which was also seen in Topology A. While acceptable predictions were made in Topology A, possibly due to the smaller ground truth elasticity and FCC values in the network, excluding fluxes in Topology B led to BMCA predicting seemingly random values that roughly matched the sign (positive or negative) of ground truth values. These predictions were neither accurate nor informative, further reinforcing the conclusion that flux data is essential for accurate FCC predictions.

**Fig 14.**
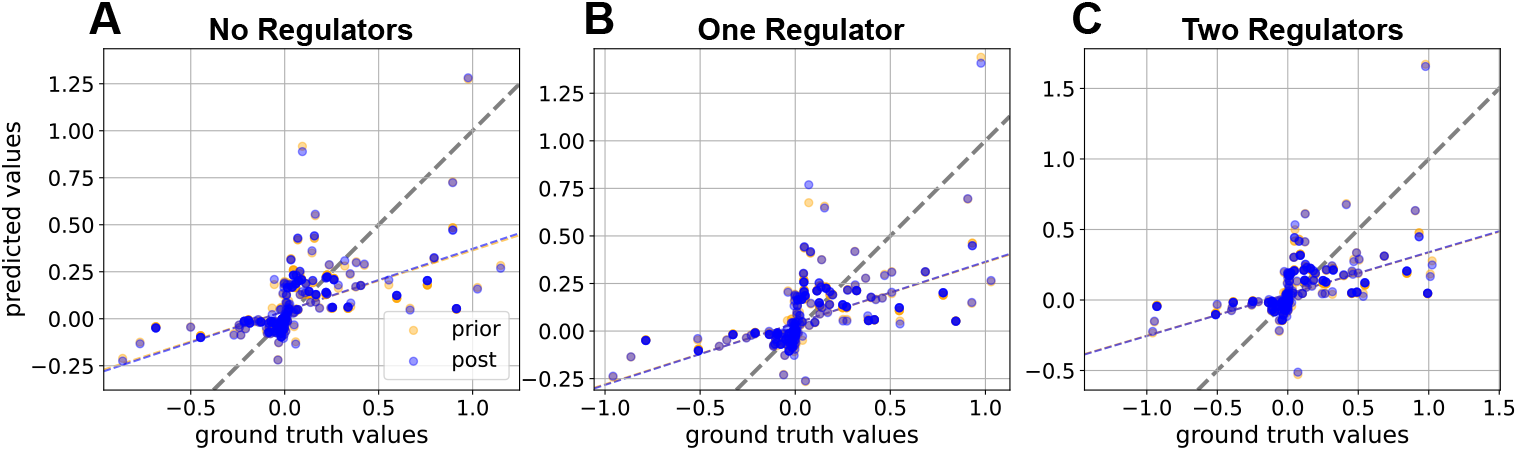
BMCA FCC predictions compared against ground truth values for Topology B model variations when flux values were omitted. A) no regulators, B) one regulator, C) two regulators. Each dot signifies the median while the error bars represent the range of the predicted elasticity values across the different enzyme perturbation levels tested. The titles for each graph indicate which data type was omitted when running BMCA. Corrections for FCC values for reactions whose enzymes are being perturbed have been applied.

In contrast, excluding internal metabolite concentration data in Topology B led to FCC predictions that were close to their respective ground truth values across all allosteric variations. While the elasticities predicted for model variations with more allosteric regulators exhibited more variance, nearly all predictions were close (within 0.5) of their ground truth values. This finding aligns with the observation in Topology A, supporting the conclusion that BMCA does not strongly rely on internal metabolite concentrations to make FCC predictions. Excluding enzyme data in Topology B resulted in many overestimated FCC predictions, a pattern that paralleled the underestimation of the elasticity values when the enzyme data was omitted. This underscores the importance of enzyme data in obtaining good predictions for both elasticities and FCC values, unlike was observed when omitting the internal metabolite concentrations. Finally, the FCCs predicted by the BMCA algorithm for TopC demonstrated an improvement over predictions based solely on prior information, with overestimations of values near zero being minimized (Fig. 15). The median FCC values across different perturbation strengths fell within the corresponding uncertainty ranges; however, the uncertainty ranges remained too large to provide meaningful insights.

**Fig 15.**
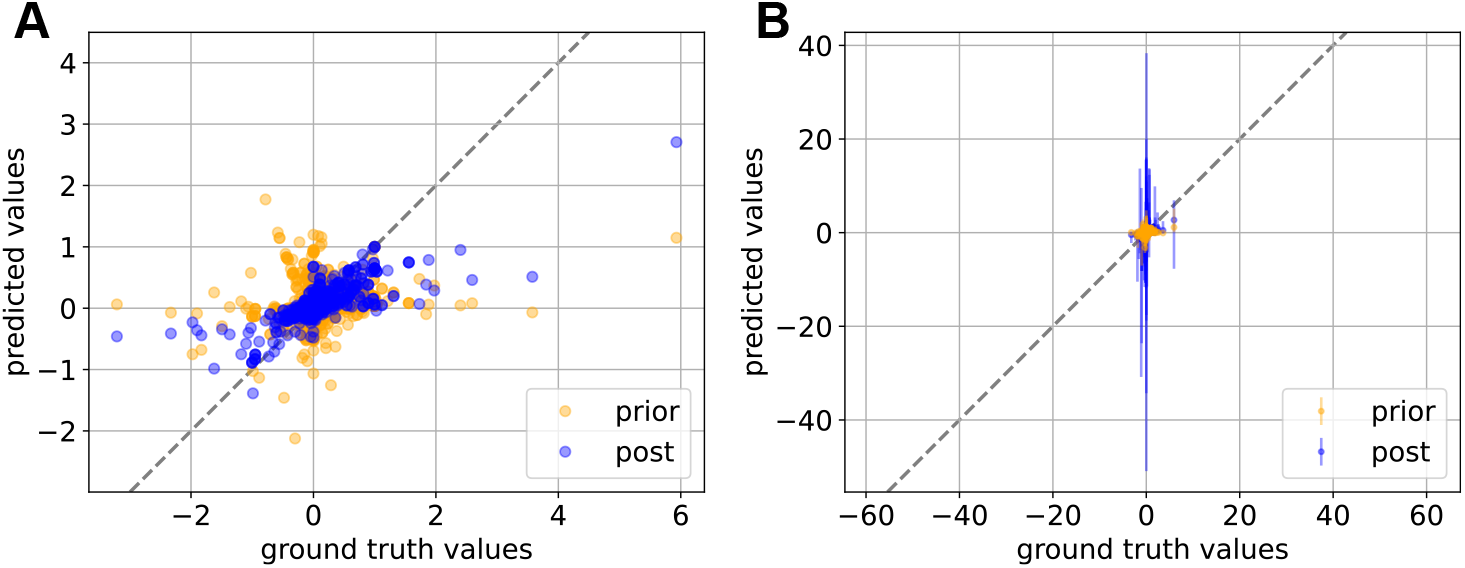
Topology C FCC value comparisons. A) Median of FCC values calculated from all perturbations of HDI elasticity distributions predicted by BMCA. B) Ranges of predicted FCC values across all perturbation strengths.

### Strength of allostery affects elasticities predicted by BMCA

Previous work [11] on BMCA demonstrated the potential for identifying latent allosteric relationships using the BMCA algorithm, albeit with limited reliability. To further investigate the BMCA algorithm’s capacity to detect allosteric regulation, reg1 and reg2 variations of synthetic models Topology A and Topology B were employed. Initially, the Hill coefficients for these allosteric regulators were set to 1, resulting in weak regulatory strength. Subsequently, the strength of the allosteric relationships was enhanced by increasing the Hill coefficients of the metabolites (Tables 4, 5). The impact of these modifications on the BMCA algorithm’s predictive accuracy was then assessed.

**Table 4.** Ground truth elasticity values before and after allostery was increased for Topology A. The first value indicates the original ground truth elasticity value and the second value indicates the ground truth elasticity value after the allosteric strength was increased.

|  | Metabolite J on reaction v6 |  | Metabolite G on reaction v3 |  |
| --- | --- | --- | --- | --- |
|  | *Original ground truth value* | *Ground truth value after increase in Hill coefficient* | *Original ground truth value* | *Ground truth value after increase in Hill coefficient* |
| TopA-reg1 | -0.680 | -3.611 | n/a |  |
| TopA-reg2 | -0.667 | -3.517 | -0.655 | -2.201 |

**Table 5.** Ground truth elasticity values before and after allostery was increased for Topology B. First value indicates the original ground truth elasticity value and the second value indicates the ground truth elasticity value after the allosteric strength was increased.

|  | Metabolite H on reaction v5 |  | Metabolite O on reaction v14 |  |
| --- | --- | --- | --- | --- |
|  | *Original ground truth value* | *Ground truth value after increase in Hill coefficient* | *Original ground truth value* | *Ground truth value after increase in Hill coefficient* |
| TopB-reg1 | -0.566 | -2.604 | n/a |  |
| TopB-reg2 | -0.566 | -2.605 | -0.439 | -3.278 |

The BMCA algorithm did not accurately predict the elasticity of allosteric regulators in both Topology A and Topology B, regardless of whether one or two regulators were present. The elasticities for these regulators were consistently predicted as zero (Fig. 16, 17). Increasing the strength of the allosteric regulators did not improve the predictiveness of the elasticities; BMCA continued to significantly underestimate the ground truth elasticity of the strengthened allosteric regulators, frequently predicting zero. However, in Topology B, the predictiveness of elasticities for non-allosteric metabolites with ground truth magnitudes larger than -2 and 2 improved, particularly when the perturbation strength was larger (Fig. 17). Specifically, in Topology B-reg1, predictions became more accurate as perturbation strength increased, though this trend was not consistently observed across other model variations.

**Fig 16.**
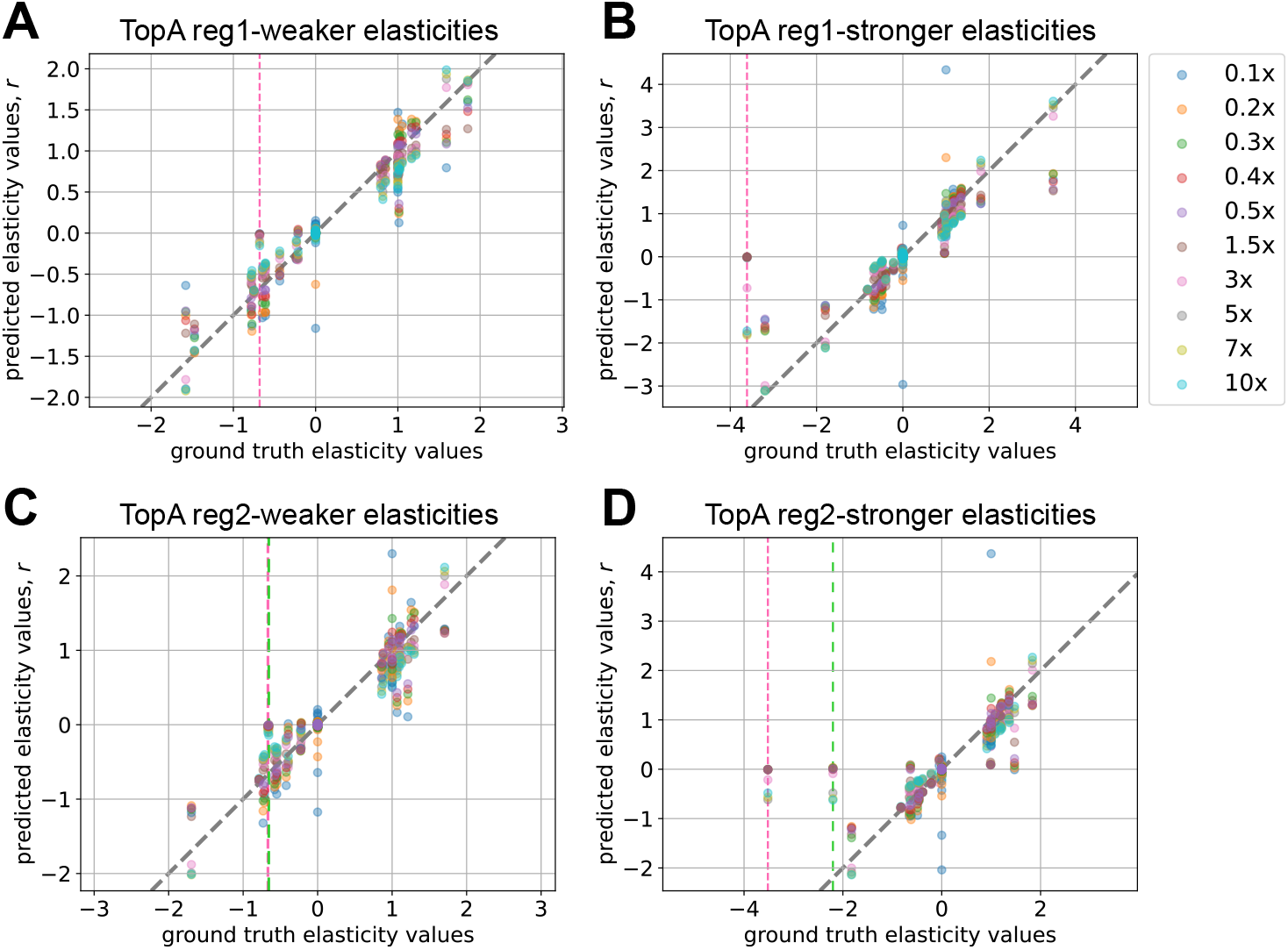
Elasticity predictions for Topology A with regulators. A) One regulator, no increase in allosteric regulation; B) One regulator, with the allostery increased; C) Two regulators, no increase in allosteric regulation; D) Two regulators, with allostery increased. The dots represent the elasticity predictions made by the BMCA algorithm using datasets with varying perturbation levels. The gray dashed line represents where the values predicted by the BMCA algorithm and the ground truth elasticity values match. The pink dashed line denotes the ground truth elasticity value of the first regulator and the green dashed line denotes the ground truth elasticity value of the second regulator.

**Fig 17.**
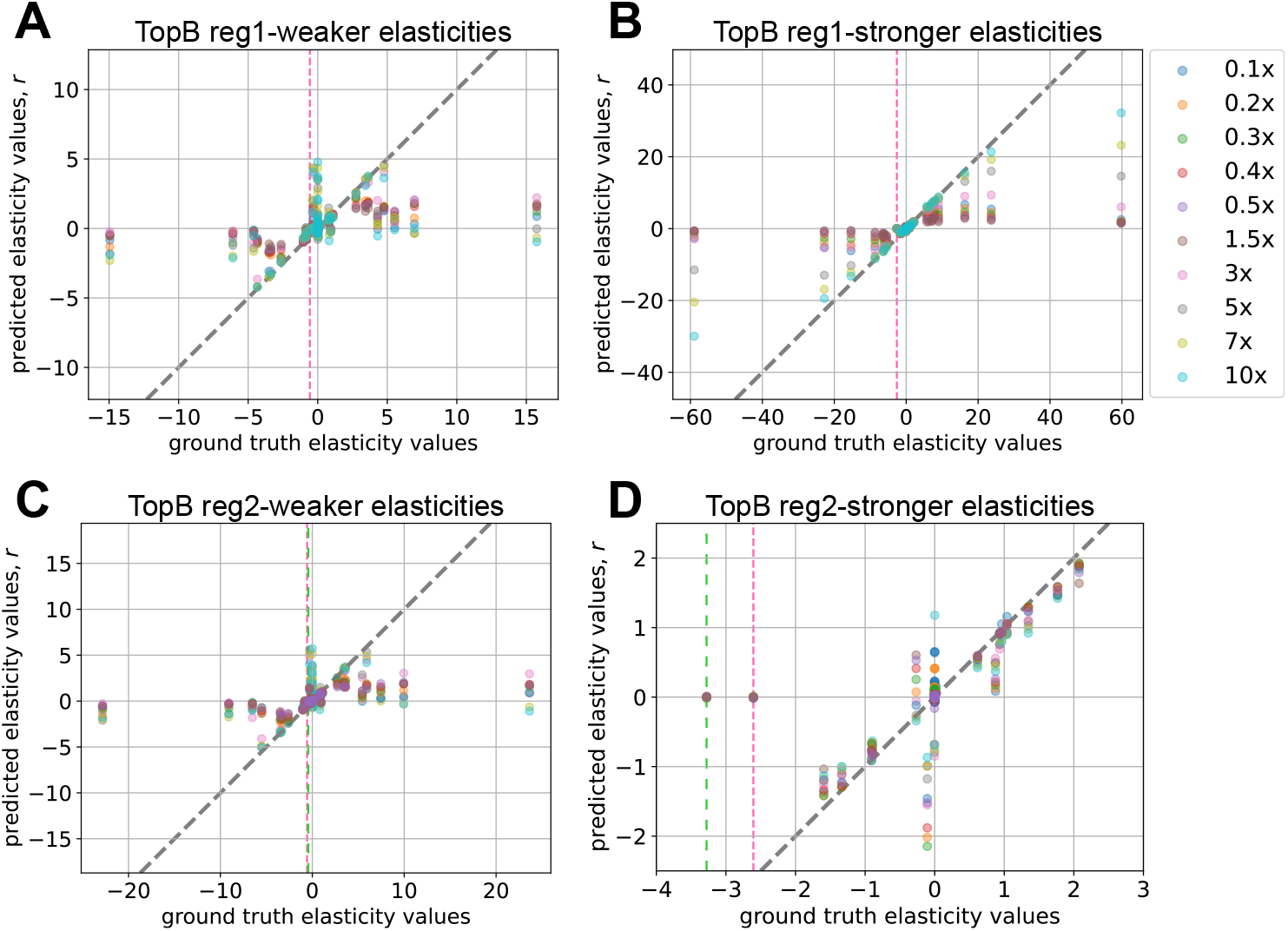
Elasticity predictions for Topology B with regulators. A) One regulator, no increase in allosteric regulation; B) One regulator, with the allostery increased; C) Two regulators, no increase in allosteric regulation; D) Two regulators, with allostery increased. The dots represent the elasticity predictions made by the BMCA algorithm using datasets with varying perturbation levels. The gray dashed line represents where the values predicted by the BMCA algorithm and the ground truth elasticity values match. The pink dashed line denotes the ground truth elasticity value of the first regulator and the green dashed line denotes the ground truth elasticity value of the second regulator.

### Evaluating FCC rankings predicted by BMCA

The informativeness of a single FCC value is limited. To determine the significance of a reaction’s FCC value, it must be compared to the FCCs of all other reactions in the network. Given the practical constraints of flux and enzyme concentration data availability, this study aimed to evaluate whether the predicted FCC values maintained their relative magnitudes in accordance with the order of ground truth FCC values. To assess the influence of different data types on the BMCA algorithm’s ability to predict FCC values and rankings, Spearman correlations were calculated between predicted and ground truth rankings across runs with varying allosteric regulators and omitted data types (Fig. 18).

**Fig 18.**
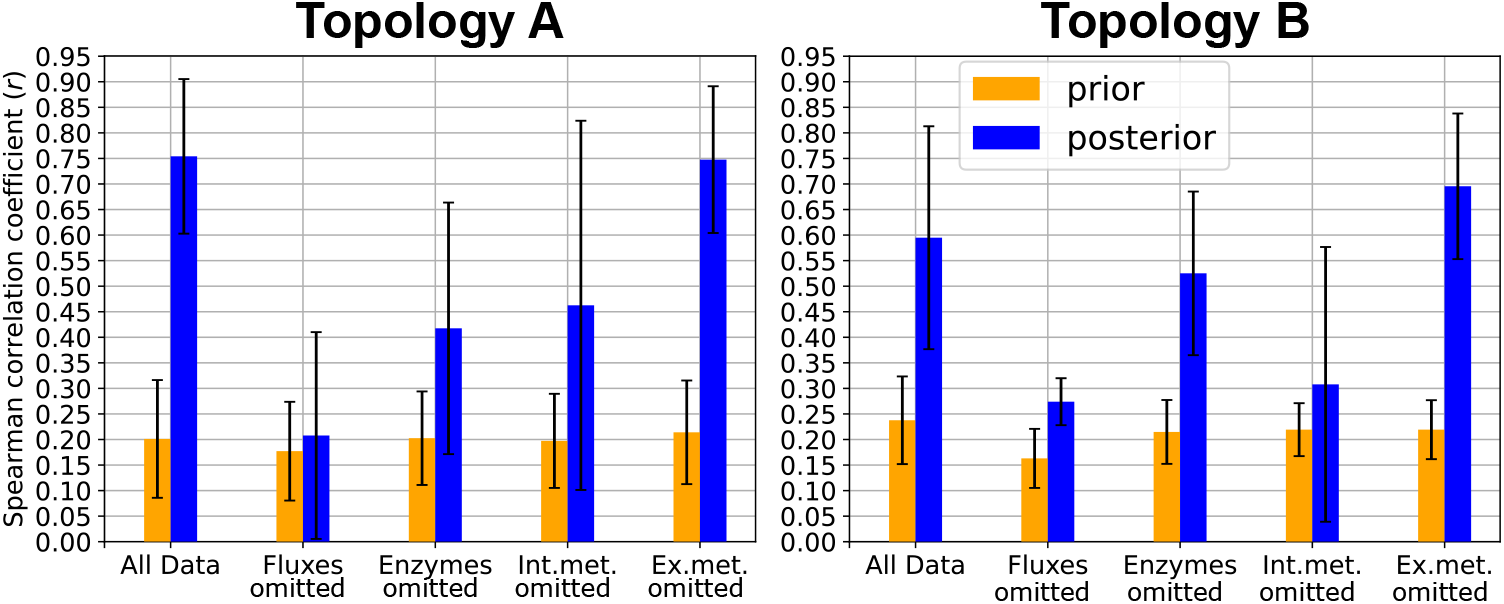
Aggregated Spearman correlation coefficients from comparisons between ground truth rankings of FCC values and FCC values predicted by the BMCA algorithm.

The difference in the change of correlations between the prior and posterior distributions were significant for many of the different data-type input trials, suggesting that the BMCA algorithm changed the relative FCC values based on the inputted data. The highest FCC values are the most useful for engineering strains, as reactions with these values represent the most promising targets. Therefore, the ten reactions with the highest predicted FCC values were examined to determine if they aligned with the order of the ground truth FCC values. Across all of the different variations of data-type inputs to the BMCA algorithm and the perturbation levels of the inputted data, the number of correctly predicted reactions predicted to be in the top ten was about the same. There was also little difference between the number of reactions that were predicted to be in the top ten calculated from the prior and posterior distributions. Only when all data was made available or the external metabolites were omitted was there a definitive improvement in FCC ranking prediction (Fig. 19). While the omission of enzyme concentration and internal metabolite concentration data resulted in minor improvements in predicting the top ten FCC enzymes, there was greater statistical overlap with prior counts. Notably, there was no change in identifying the top ten ranked FCCs when flux values were omitted between the prior and posterior predictions. This pattern persisted even in the presence of allosteric regulation S2 Fig. Similarly, for Topology C, there was a marginal increase between using the prior or posterior distribution to calculate the number of correctly identified reactions with the highest FCCs when all data types were provided to BMCA (Fig 19). BMCA is able to predict at least 4 of the top 10 reactions with the highest FCC rankings for Topology C, and inputting all possible data into the BMCA algorithm only improved the prediction of the top ten FCCs by one more metabolite.

**Fig 19.**
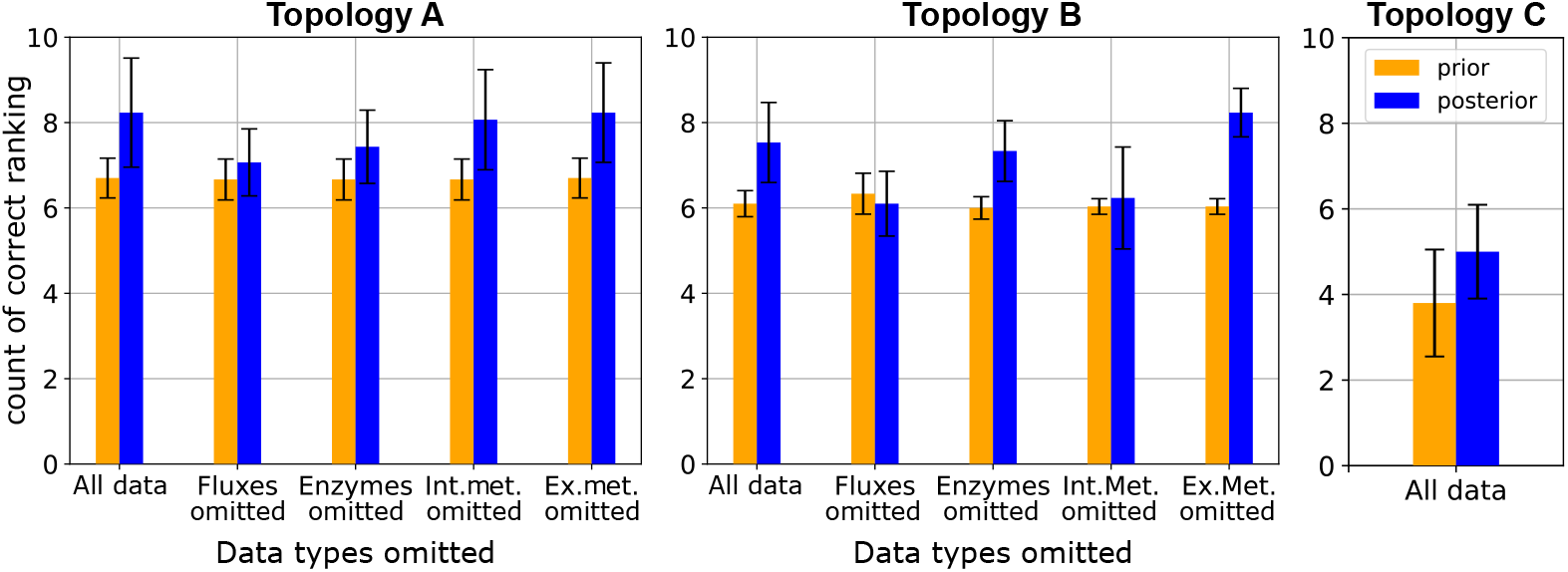
Number of enzymes correctly predicted as having one of the top ten highest FCC values based on the various data types omitted when running BMCA. The height of the bars represent the mean of the predictions across all the perturbation levels and allosteric regulation levels; the error bars represent the standard deviation.

Analysis of all FCC values predicted by BMCA for Topology C reveals that the ranking of FCC values is predominantly influenced by the observed reaction rather than the perturbed enzyme. This observation is supported by the distinct vertical patterns evident in the heatmap (Fig. 20). These patterns suggest that the underlying network structure exerts a significant influence on FCC values, potentially overshadowing the direct effects of individual enzyme perturbations.

**Fig 20.**
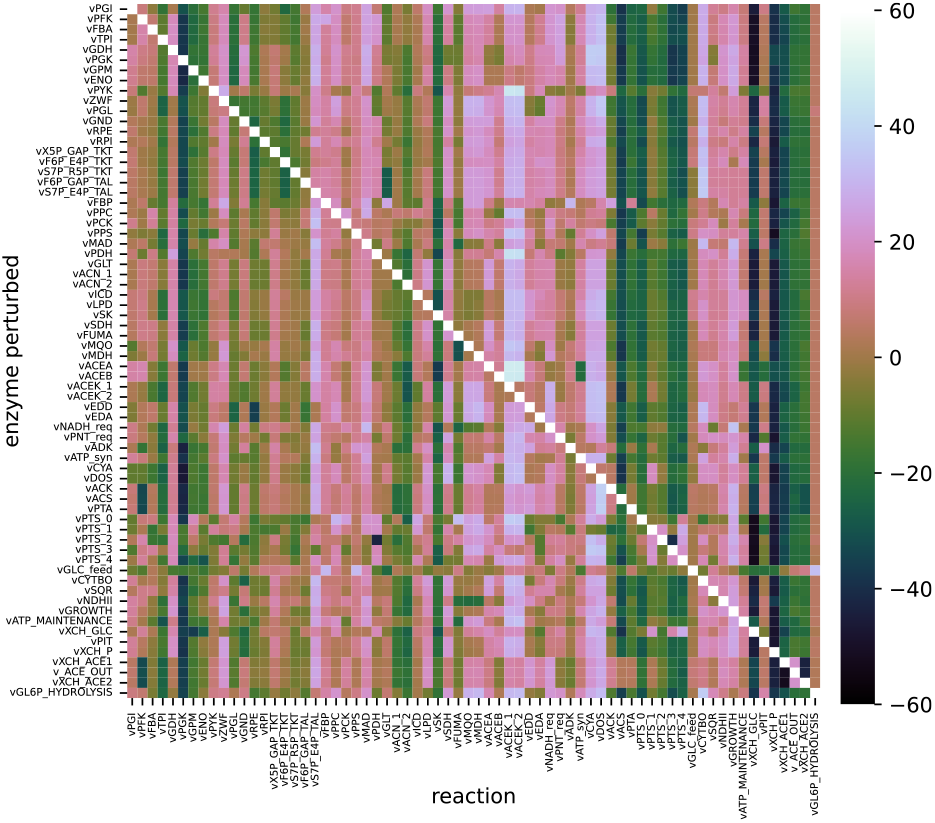
Accuracy of FCC rankings predicted by the BMCA algorithm. Y-axis describes which enzymes were perturbed while the x-axis describes the FCC being observed. The colorbar quantifies the difference in predicted ranking from the ground truth ranking.

The true rankings of the enzymes that the BMCA algorithm incorrectly identified as being in the top ten were further investigated across the ten different perturbation levels tested on Topology C without any omissions of any data (Fig. 21). The BMCA algorithm most frequently predicted that “XCH ACE1,” the reaction transporting acetate out of the cell into the periplasm, and “XCH P,” the reaction transporting phosphate into the cell from the periplasm, were in the top ten. However, these reactions’ ground truth FCC rankings were 57 and 61st, respectively.

**Fig 21.**
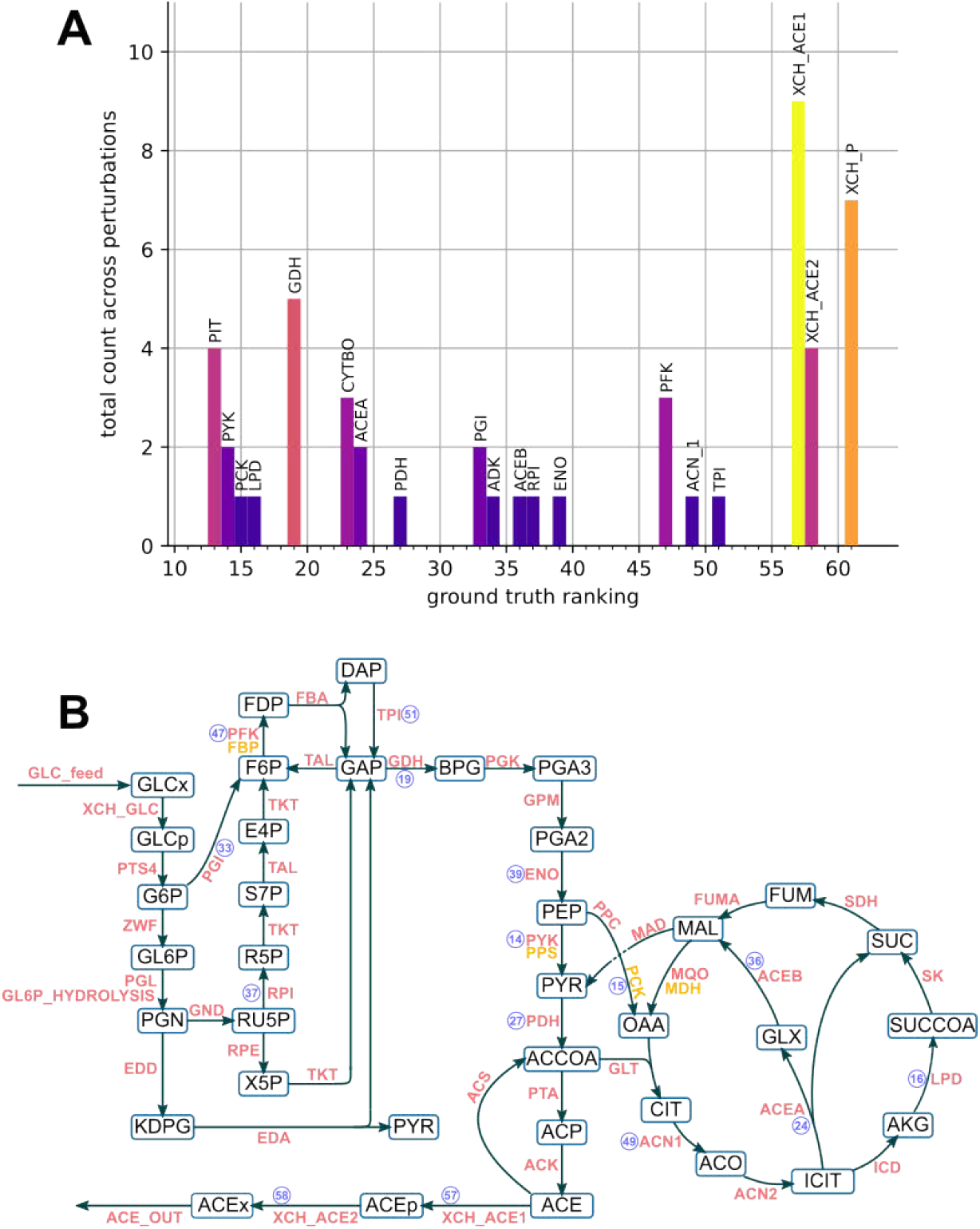
Ground truth rankings of enzymes that were incorrectly predicted as having a top ten highest FCC value for Topology C. A) The bars represent the number of times an enzyme with a non-top ten ground truth ranking was incorrectly predicted across a total of ten different perturbations. B) Ground truth rankings shown in purple next to the reactions that were incorrectly predicted as being in the top 10. Reactions consisting of cofactors only are not shown.

## Conclusion

Our study aimed to evaluate the predictive capabilities and limitations of BMCA in determining key control points within metabolic pathways, particularly in data-limited scenarios. By systematically assessing the impact of different physiological data types on the BMCA algorithm’s accuracy at predicting control coefficients and allosteric interactions with our three synthetic models, we provide a clearer understanding of its strengths and constraints. Overall, given sufficient data, we demonstrate that BMCA can make reasonable predictions of control coefficient values and relative orders.

Additionally, we show that allosteric regulations latent in the data are not predicted well by the BMCA algorithm. This work contributes to the broader effort of optimizing metabolic engineering by offering guidance on when and how BMCA can be effectively applied. More generally, our findings underscore the potential of mechanistic models to provide biological insight while requiring less extensive data collection compared to machine learning approaches, positioning them as valuable tools for accelerating the rational design of biomanufacturing processes.

### Impact of Data Availability on BMCA Predictive Accuracy

An important consideration in the application of BMCA is its robustness to missing data, given the challenges associated with obtaining experimental measurements.

Understanding the relative importance of different data types can provide valuable guidance for prioritizing data collection efforts. The impact of omitting specific data types was assessed across different network topologies, revealing distinct effects on model accuracy (Table 6).

**Table 6.** Summary of effects on different control coefficients when different types of data types are omitted from BMCA.

| Data Omitted | Elasticities | CCCs | FCCs |
| --- | --- | --- | --- |
| None | Predictive | Predictive, with some poor estimation | Good, with some poor estimation |
| Fluxes | Signs only | Uninformative | Uninformative |
| Enzyme Concentrations | Underestimation | Poor estimation | Poor estimation |
| Internal Metabolites | Underestimation | Poor estimation | Predictive |
| External Metabolites | Predictive | Predictive | Predictive |

Omitting external metabolite data had contrasting effects on the predictive accuracy of the BMCA algorithm for different topologies. In Topology A, this omission had minimal impact on elasticity, CCC, and FCC predictions, suggesting that external metabolite concentrations contribute little to the algorithm’s performance in smaller systems. However, for Topology B, excluding external metabolites improved predictions across all metrics. Internal metabolite data played a significant role in elasticity and CCC predictions. Without this data, elasticity values were strongly underestimated, often near zero. Since CCCs are derived from elasticities, they were subsequently poorly estimated. Interestingly, FCC predictions remained accurate despite the omission, as the overestimation of CCCs and underestimation of elasticities effectively canceled out when calculating FCCs. The omission of enzyme data had a more pronounced effect. Elasticity predictions were underestimated, often nearing zero, suggesting that enzyme values provide critical scaling information. Without enzyme data, both CCC and FCC values were poorly estimated. Unlike internal metabolite data, the lack of enzyme information appears to create a scaling issue that persists through the FCC calculations, as the scaling factor was missing in the elasticity-to-CCC transformation. Flux data was crucial for accurate predictions across all metrics in both topologies. Excluding flux data degraded the quality of elasticity, CCC, and FCC predictions, preserving only the sign of the ground truth values while failing to capture their magnitudes. The elasticity and FCC predictions also demonstrated the crucial role flux data plays in informing the BMCA algorithm’s predictions. There was no difference between the prior and posterior distributions without the flux data, even with the presence of all other types of data. Flux data, like enzyme data, plays a key role in scaling elasticities during CCC calculations. Its absence leads to compounded inaccuracies, making FCC predictions particularly unreliable. In conclusion, external metabolite data is largely non-critical for BMCA predictions and may even be detrimental in some systems. By contrast, flux and enzyme data are essential for accurate predictions, with their omission causing significant errors. Internal metabolite data strongly influences elasticity and CCC predictions but has limited impact on FCC accuracy. Finally, network complexity amplifies the effects of data omissions, especially for flux and enzyme data, as observed in Topology B.

### Predicting Allosteric Regulators

While the ability to predict allosteric regulation based on latent data patterns is a promising aspect of BMCA, the algorithm repeatedly failed to identify allosteric regulation across multiple models, even when the regulatory effects were strong. This raises the question of what specific conditions are required for BMCA to detect allostery, particularly since previous studies (11) identified only a single allosteric interaction in a medium sized model or identified the interaction in a model with only three steps and three metabolites. Additionally, the presence of a single high-value allosteric elasticity in the system appeared to allow BMCA to overcome its typical constraint of predicting elasticity values within the range of -1.5 to 1.5, suggesting that larger perturbations may enhance predictive accuracy. This outcome is unexpected because, despite the presence of several non-allosteric elasticities with higher magnitudes in Topology C, their predicted values remain constrained within the 1.5 magnitude limit. Although predictions remained inaccurate even when allosteric regulation strength was high, the overall accuracy of elasticity value predictions improved, indicating that BMCA extracts meaningful relationships between metabolites from the data but is unable to explicitly attribute these relationships to specific allosteric interactions.

### Impact of Network Structure on FCC Ranking Fidelity

While absolute FCC values can provide useful insights when considered in the context of the entire system, an alternative approach is to rank all FCC values by magnitude to identify the most influential reactions. To evaluate the ranking fidelity of the BMCA algorithm, its ability to correctly identify the top-ranking FCC values was assessed. The results indicated that providing additional data led to only limited improvements in the accuracy of the top ten FCC predictions. Furthermore, predicted FCC rankings remained largely unchanged regardless of which enzyme was perturbed during data collection. These findings suggest that network structure plays a fundamental role in determining FCC values within BMCA, highlighting the need for further investigation into its influence on metabolic control predictions.

### Elasticity limitations

Understanding metabolic network behavior requires accurate predictions of elasticity values, which describe how reaction rates respond to changes in metabolite concentrations. The BMCA algorithm was developed to estimate these values, but its accuracy varies depending on reaction topology and elasticity range. While it performs well for reactions with small elasticity values, it struggles with larger elasticities, particularly outside the range of -1.5 to 1.5. In these cases, the algorithm systematically underestimates values, capping predictions around ±2. This limitation is especially pronounced for reactions near equilibrium, where elasticity magnitudes of 10 or greater are often reduced to 1.5 or lower. These findings suggest that BMCA is more reliable for moderate elasticity ranges but requires improvement in capturing extreme values. To enhance BMCA’s predictive utility, it is essential to determine an appropriate range for elasticity values. One way to do this is by analyzing the disequilibrium ratio (*ρ*), which quantifies how far a reaction is from equilibrium. Since reactions near equilibrium contribute minimally to metabolic control, accurately predicting their elasticity coefficients is less critical. Instead, it is more important for the BMCA algorithm to prioritize identifying reactions further from equilibrium, which are more relevant for strain design. The disequilibrium ratio is defined as the mass-action ratio divided by the equilibrium constant (equation 1, 2), where the product and substrate concentrations determine the deviation from equilibrium. When *ρ* = 1, the reaction is at equilibrium. When *ρ <* 1, the reaction is out of equilibrium in the forward direction. Likewise, when *ρ >* 1, the reaction is out of equilibrium in the reverse direction. The magnitude of elasticity values depends on the disequilibrium value (Equation 3). Large-magnitude elasticities typically correspond to *ρ* values close to 1, signifying near-equilibrium conditions, while small-magnitude elasticities indicate reactions further from equilibrium (17).

BMCA’s current elasticity predictions, where magnitudes of elasticities are capped at 2, constrain the predicted disequilibrium ratio to between 0.5 and 1.5 in the forward and reverse direction, respectively. However, reactions with elasticity magnitudes greater than 2 can still be far from equilibrium, meaning this cap leads to underestimated elasticities and, consequently, overestimated concentration and flux control coefficients.

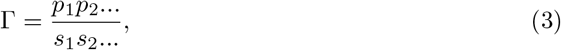

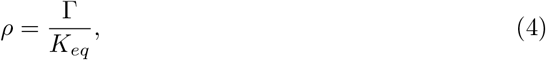

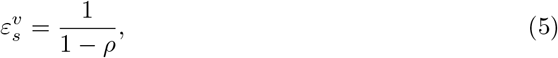

### Effect of perturbation and allostery levels on BMCA predictions

Perturbation levels appear to have little effect on TopA, possibly due to its smaller elasticities, which may facilitate accurate predictions even in the presence of allosteric regulators. In contrast, for TopB and TopC, perturbation levels influence elasticity predictions, particularly when allostery is introduced. The extent of allostery does not appear to affect the predictive performance of the BMCA algorithm; rather, perturbation level plays a more significant role. However, determining which perturbation levels yield more accurate predictions and understanding the underlying reasons remain challenging. Further investigation is needed to address these questions.

## Future work

At present time, BMCA only provides small improvements in estimating elasticities. Further investigation is needed to validate these findings under different conditions. Future research could add noise in the data tested, employ different prior distributions, and investigate the combination of flux and internal metabolite concentrations necessary for BMCA to provide a more comprehensive understanding of the limitations of BMCA. Refining the Bayesian framework of BMCA may enhance its predictive power. The influence of prior distributions remains an open question, as current results exhibit a strong bias toward predicting zero for all elasticities and a tendency to constrain estimates within the range of -1.5 to 1.5. Testing alternative prior distributions that would expand the BMCA algorithm’s predictive range for elasticities from -5 to 5 could improve its estimation accuracy and better capture the elasticity values whose substrate disequilibrium ratios fall within 1± 0.2. Additionally, introducing controlled levels of noise to the ground truth data would provide insight into BMCA’s robustness under realistic experimental conditions, where measurement variability is inevitable. BMCA currently requires steady-state data, verified by the multiplication of the stoichiometric matrix (N) with the reference fluxes (v*), but assessing its sensitivity to deviations from steady-state assumptions could further refine its applicability. BMCA’s inability to detect allosteric regulation remains a key limitation. Future work should aim to identify the specific conditions under which BMCA can successfully infer allosteric interactions. Investigating whether strong perturbations or particular network configurations influence allosteric detection could provide insight into the algorithm’s constraints. By addressing these areas, future research can refine BMCA’s predictive capabilities, improve its ability to infer metabolic control mechanisms, and clarify the conditions under which it is most effective.

## Data and code availability

The data generated and analyzed during the current study are available in the GitHub repository: https://github.com/sys-bio/BMCA-pipeline

## Supporting information

**S1 Fig.**
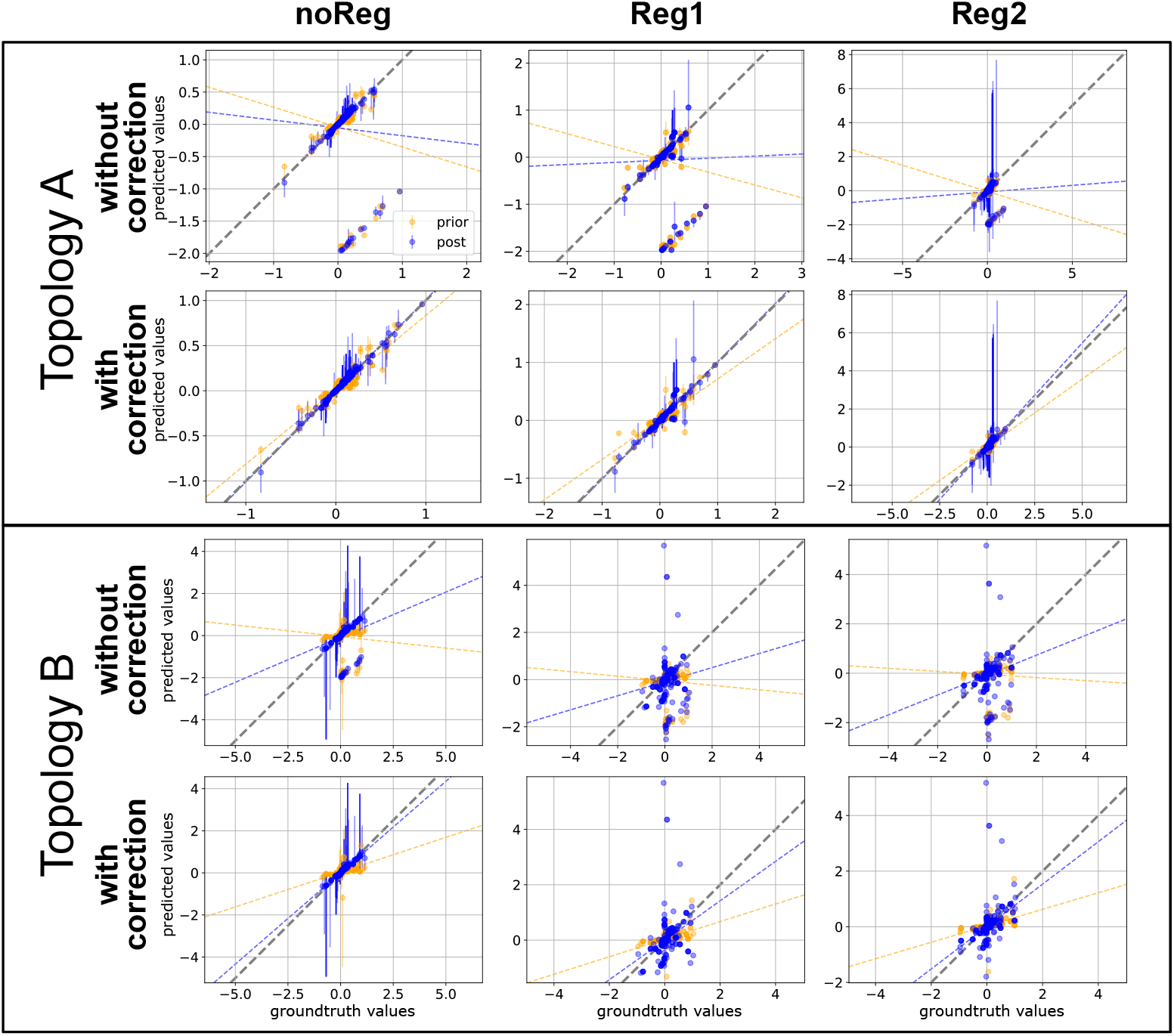
Difference in corrected and uncorrected FCC predictions. Each dot signifies the median of the predicted values across the different enzyme perturbation levels tested. The grey dashed line represents where the ground truth values match the predicted values. The other dashed lines represent linear regression for the dots of the same color.

**S2 Fig.**
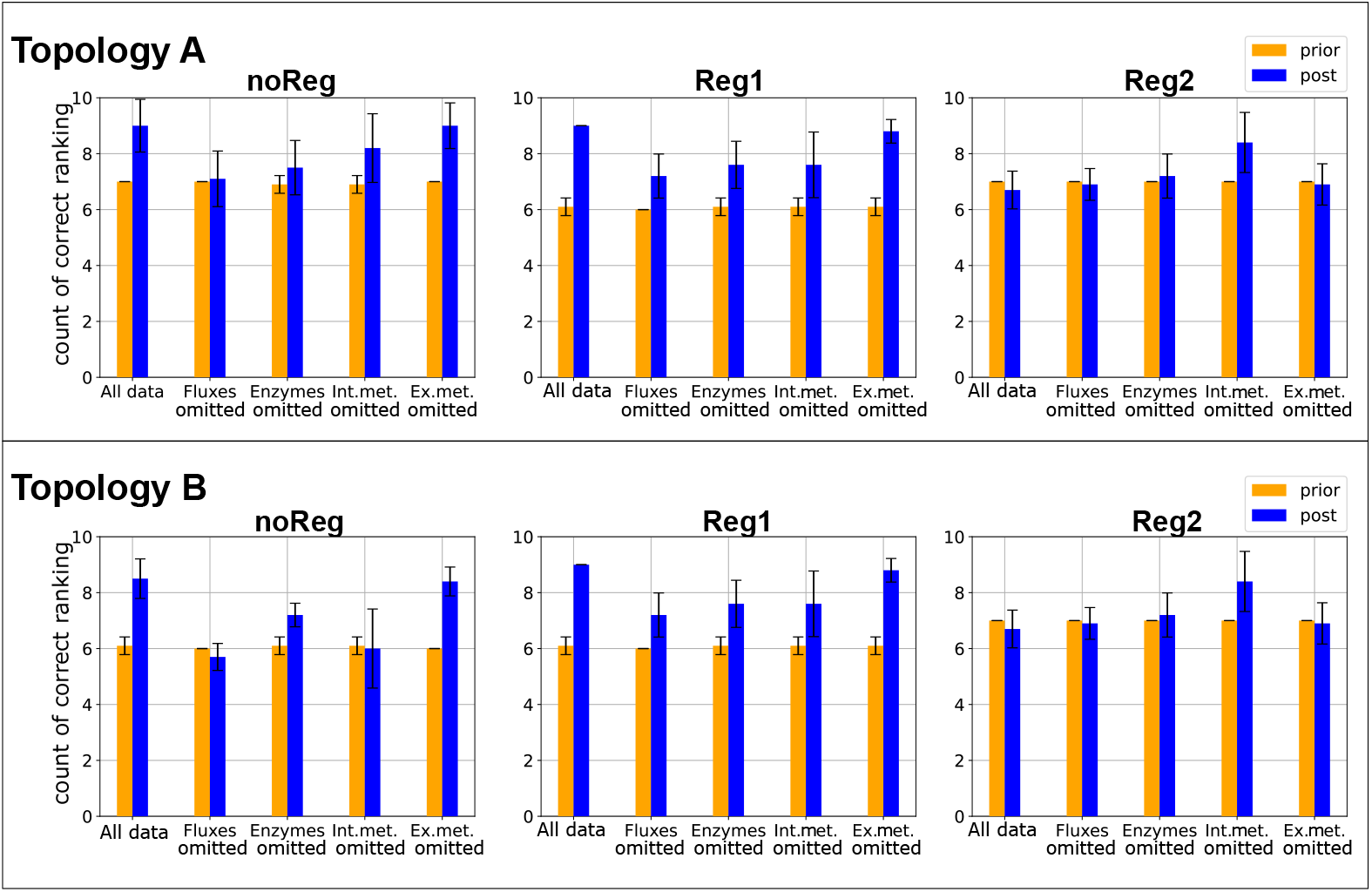
Number of enzymes correctly predicted as having one of the top ten highest FCC values based on the various data types omitted when running BMCA across different amounts of regulators. The height of the bars represent the mean of the predictions across all the perturbation levels and allosteric regulation levels; the error bars represent the standard deviation.

## Acknowledgments

We thank Peter St. John, Jeremy Zucker, and Andrew McNaughton for helpful discussions.

